# An analogue of the Prolactin Releasing Peptide reduces obesity and promotes adult neurogenesis

**DOI:** 10.1101/2023.02.28.528936

**Authors:** Sara K. M. Jörgensen, Alena Karnošová, Simone Mazzaferro, Oliver Rowley, Hsiao-Jou Cortina Chen, Sarah J. Robbins, Sarah Christofides, Florian T. Merkle, Lenka Maletínská, David Petrik

## Abstract

Hypothalamic Adult Neurogenesis (hAN) has been implicated in regulating energy homeostasis. Adult-generated neurons and adult Neural Stem Cells (aNSCs) in the hypothalamus control food intake and body weight. Conversely, Diet Induced Obesity (DIO) by High Fat Diets (HFD) exerts adverse influence on hAN. However, the effects of anti-obesity compounds on hAN are not known. To address this, we administered a lipidized analogue of an anti-obesity neuropeptide, Prolactin Releasing Peptide (PrRP), so-called LiPR. In the HFD context, LiPR rescued survival of adult-born hypothalamic neurons and increased the number of aNSCs by reducing their activation. In addition, LiPR rescued reduction of immature hippocampal neurons and modulated calcium dynamics in iPSC-derived human neurons. These results show for the first time that anti-obesity neuropeptides influence adult neurogenesis and suggest that the neurogenic process can serve as a target of anti-obesity pharmacotherapy.

## Introduction

Energy homeostasis is regulated in the hypothalamus, a ventral part of diencephalon critical for basic physiological functions.^1,2^ In the Medial Basal Hypothalamus (MBH), anorexigenic (i.e., food-intake inhibiting) and orexigenic (i.e., food-intake promoting) neurons sense metabolites and endocrine signals from the periphery to regulate eating behavior and appetite.^3,4^ Anti-obesity compounds lower the sensation of appetite and the body weight by targeting neurons in the MBH.^5^ Unfortunately, several anti-obesity compounds originally approved for clinical use, were later rejected for severe side effects such as fetal toxicity, nausea, or depression.^6–8^ The only neuroactive medicines with anti-obesity activity approved in the UK and EU are analogues of the GLP-1RA such as Liraglutide (Saxenda) or Semaglutide (Ozempic).^9–11^ However, their primary indication is for Type-2 diabetes mellitus (T2DM).^12–14^ Clearly, there is a need for novel neuroactive anti-obesity compounds. Lipidized analogues of PrRP are a novel class of compounds with robust anorexigenic and anti-obesity effects.^15,16^

Despite its name, PrRP does not regulate prolactin release ^17^, but is the Prlh gene-encoded anorexigenic neuropeptide ^18^, released by neurons in the hypothalamus ^19^, which increases neuronal survival and reduces food intake. ^20–23^ However, PrRP is Blood Brain Barrier (BBB)-impermeable, and its activity is limited only to minutes. ^23^ To circumvent these limitations, a lipidized analogue of PrRP was developed, which has extended pharmacokinetics and can exert its effect in the Central Nervous System after peripheral administration.^15^ This full-length, 31 amino acid long PrRP peptide palmitoylated on Lysine-11 (palm^11^-PrRP31), further referred as LiPR, activates cells in MBH to reduce food intake, fat mass and weight gain (by around -15%) in mice exposed to the obesity-inducing High Fat Diet (HFD).^16,24^ To exert its anorexigenic effects, LiPR binds to PrRP transmembrane G-protein Coupled Receptors, GPR10 and NPFF2 ^19,25,26^ with higher affinity than PrRP and 2-times higher affinity for NPFF2 than GPR10. ^24,27^

While many of the appetite-controlling neurons in the MBH originate in embryogenesis, some of them are newly generated during adulthood from adult Neural Stem Cells (aNSCs), which reside in the Hypothalamic Ventricular Zone (HVZ) in the process of hypothalamic adult neurogenesis (hAN). ^28,29^ Both hypothalamic aNSCs and newborn MBH neurons are critical for energy homeostasis. ^30,31^

In HVZ, tanycytes, radial-glia-like cells that line the walls of 3^rd^ ventricle (3V), serve in a dual role as the putative hypothalamic aNSCs (htNSCs) ^30,32^ and as metabolic regulators ^33^ to control energy homeostasis, body weight ^34–36^ and access of anti-obesity compounds to the brain parenchyma. ^37^ Tanycytes give rise to differentiated cell progeny such as glial cells or neurons ^30,38,39^, which are part of almost 2000 cells generated daily in the adult murine hypothalamus.^31^ New hypothalamic neurons display unique properties different from embryo-generated neurons ^40^ and make up to 37% of all hypothalamic neurons. ^41^

DIO caused by the HFD has been shown to reduce hAN through lower survival and higher apoptosis of newborn neurons or by decreased proliferation of tanycytes.^34,35,42^ On the other hand, ablation of hAN leads to higher weight gains in HFD-fed animals ^30,31^, whereas increasing hAN protects against adverse effects of HFD ^30^ suggesting that newborn neurons are anorexigenic. However, the relationship between hAN and DIO is insufficiently understood and marred by conflicting results. ^34,35,42^

Given the importance of hAN for energy homeostasis, we hypothesized that anti-obesity compounds may be neurogenic in a similar manner to anti-depressants. ^43,44^ To test this hypothesis, we administered LiPR and Liraglutide to mice exposed to HFD and determined the effects of this treatment on cellular and molecular processes of hAN. Our results show that HFD reduces number of htNSCs in HVZ and survival of newborn neurons in MBH. In addition, HFD downregulates adult neurogenesis in another neurogenic niche, the Subgranular Zone (SGZ) of the hippocampus ^28^. Administration of LiPR, but not Liraglutide, rescues these adverse effects of HFD on differentiation of newborn neurons in SGZ and on survival of newborn neurons and on number of hypothalamic htNSCs in MBH. LiPR also reduces proliferation and activation of hypothalamic htNSCs. This suggests that LiPR upregulates hAN by preserving the population of htNSCs and survival of new neurons. For the first time, our data show that anti-obesity treatment upregulates adult neurogenesis, which was diminished by obesity.

## Results

### LiPR lowers body weight, ameliorates metabolic parameters related to obesity and increases expression of GPR10 in MBH

To determine how HFD and LiPR affect hAN *in vivo*, adult male C57BL/6J mice were exposed to three different protocols lasting 7 days (7d; Figure 1A, Fig.1A), 21d (Fig.1B) or 4 months (4mo; Fig.1C). The short (7d) and intermediate (21d) protocols were expected to initiate HFD-induced inflammation and astrogliosis in the hypothalamus ^45,46^, whereas the long (4mo) protocol leads to DIO. ^16^ In the 7d and 21d protocols, LiPR was administered concurrently with HFD, and during the last 2 weeks of the 4mo protocol. To trace adult-generated cells, mice were administered 5-bromo-2’-deoxyuridine (BrdU) with different post-BrdU (so-called “chase”) periods as indicated in the protocol schematics (Fig.1A-C). Exposure to HFD or LiPR administration (HFD+LiPR) for 7d did not significantly change the body weight (Fig.1A). In the 21d protocol, treatment had no effect on the body weight, however, duration of treatment (two-way (TW) repeated measure (RM) ANOVA, F(9, 108) = 13.67, p < 0.0001) and interaction between treatment and its duration (F(8, 108) = 6.55, p < 0.0001) had a statistically significant effect on the body weight (Fig.1B). In contrast, there was a statistically extremely significant effect of the treatment (TW RM ANOVA, F(2, 126) = 141.3, p < 0.0001), the duration of treatment ((F(6, 126) = 7.87, p < 0.0001) and their interaction ((F(12, 126) = 34.84, p < 0.0001) on body weight in the last 2 weeks of 4mo exposure to HFD (Fig.1C). Bonferroni’s Test for multiple comparisons found that mean values of body weight were significantly higher in HFD-exposed mice than in Controls (p < 0.001) and significantly lower in HFD+LiPR mice compared with HFD (p < 0.05 at day 5 of LiPR treatment, p < 0.001 for day 7 to 14 of LiPR treatment), demonstrating anorexigenic effects of LiPR as shown before.^15^ To characterize the systemic effects of DIO, selected hormones and metabolites were measured in mouse plasma. Exposure to 4mo HFD significantly increased concentration of insulin, leptin and cholesterol but not triglycerides (TAG) or free fatty acids (FFA; Fig.1D-H), which is consistent with the development of the metabolic syndrome.^47^ Administration of LiPR for 2 weeks (wk) reduced the HFD-induced plasma concentration of insulin, leptin, and cholesterol. To exert its effects, LiPR binds to PrRP receptors GPR10 and NPFFR2 ^15^, which were identified in the hypothalamus but not in specific hypothalamic nuclei or associated with neuronal structures.^48,49^ Using immunohistochemistry (IHC), we identified GPR10-positive (GPR10+) puncta co-localizing with structures positive for the Microtubule-associated protein 2 (Map2) that labels neuronal cytoskeleton and/or surrounding neuronal nuclei expressing the marker of mature neurons, the Human neuronal protein C and D (HuC/D) ^29^, in the MBH (Fig.1I-K). This suggests that the receptor is expressed in neurons. Interestingly, exposure to 21d HFD reduced density of GPR10, which was rescued by LiPR administration (Fig.1L). In addition, over 90% of Map2+ structures associated with BrdU+ nuclei in the MBH (16d post BrdU) displayed co-localization with GPR10 puncta (Fig.1M-N) suggesting that majority of adult-generated hypothalamic neurons express this PrRP receptor. Besides GPR10, we co-localized neuronal cytoskeleton structures with NPFFR2 in the MBH (Fig.1O-P), which corresponds to its previously described expression in the hypothalamus. ^50^ Finally, we detected mRNA for PrRP and GPR10 in MBH and observed that treatment has a statistically significantly effect on their expression (Fig.1Q; TW ANOVA (TWA), F(2, 12) = 5.95, p = 0.016) with Bonferroni’s Test revealing that HFD+LiPR increases expression of Prlh compared to both Control and HFD. Taken together, these results suggest that LiPR reverses the negative effects of HFD on plasma levels of insulin, leptin and cholesterol, and its receptors localize with adult-generated cells in the MBH.

**Figure 1.**
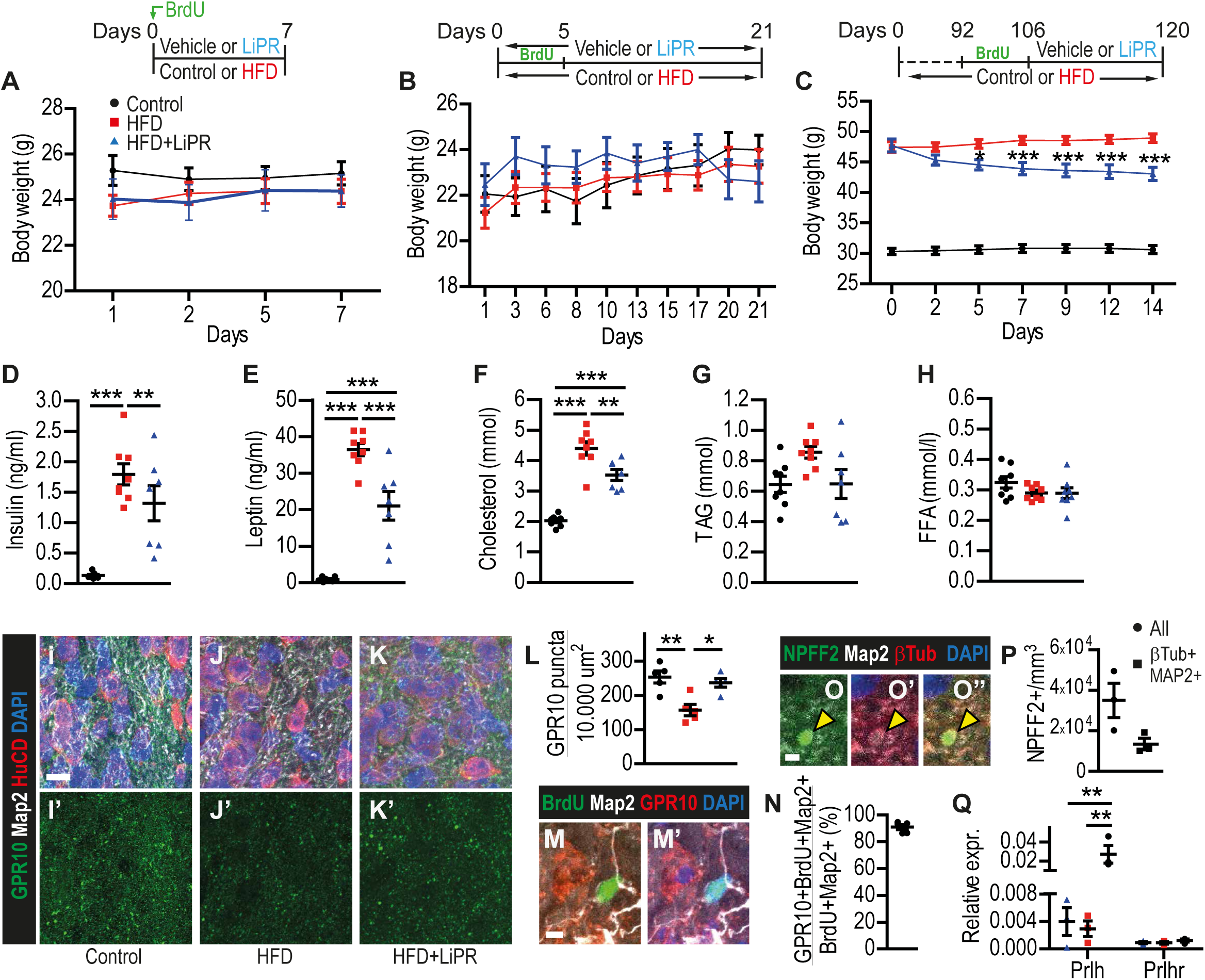
(A-C) Mouse body weigh over time for the 7d (A), 21d (B) and 4mo (C) HFD protocols, which are summarized in the schematics above panels. (D-H) Plasma concentration of selected hormones and metabolites from 4mo HFD group. (I-K) Representative confocal microphotographs of MBH stained as indicated for GPR10 and neuronal markers from 21d HFD group in Control (I), HFD (J) and HFD + LiPR (K) mice. (L) Quantification of GPR10+ puncta in MBH of 21d group. (M) A representative image of GPR10+ puncta associated with Map2+ processes around a BrdU+ nucleus in ME. (N) Quantification of the proportion of BrdU+Map2+GPR10+ cells in MBH. (O) A representative confocal image of a NPFF2 punctum in MBH stained as indicated from a control animal of 21d HFD group. (P) Quantification of NPFF2 puncta in MBH of 21d group. (Q) Relative mRNA fold change of Prlh and Prlhr compared to Gapdh in cDNA from MBH from mice 2 weeks on Control diet, HFD or HFD+LiPR. Scale bars (s.b.): 10 μm (I-K), 5 μm (M), 2 μm (O). n = 5 for 7d and 21d groups, n = 8 for 4mo group, n = 3 for neurospheres. Two-way (A-C) or one-way (all other) ANOVA with Bonferroni’s test. *p < 0.05, **p < 0.01, ***p < 0.001. Data are presented as mean ± SEM.

### LiPR rescues HFD-induced decrease in number of htNSCs

Next, we determined whether LiPR treatment influenced htNSCs (Fig.2A-C). Treatment (TWA, F(2,45) = 3.28, p = 0.047), duration of treatment (F(2,45) = 3.76, p = 0.031), and their interaction (F(4,45) = 5.29, p = 0.0014) had a statistically very significant effect on the number of GFAP+ α-tanycytes, which are considered the putative htNSCs ^51^. Bonferroni’s Test revealed a statistically significant increase in the number of GFAP+ α-tanycytes in HFD+LiPR compared to Control and HFD treatment in 7d (p < 0.05, p < 0.001, respectively), suggesting that both diet and LiPR affect number of htNSCs in the short protocol (Fig.2D). Finally, there were significantly more GFAP+ α versus 21d or 4mo HFD+LiPR groups (p < 0.001). This suggests that short LiPR rescues HFD-induced reduction in the number of htNSCs. While the absolute number of all GFAP+ tanycytes was not changed in any protocol between any groups (data not shown), there was a statistically significant effect of the treatment (TW RM F(2, 44) = 3.58, p = 0.036) and the interaction of treatment and its duration ((F(4, 44) = 3.88, p < 0.009) on proportion of GFAP+ α-tanycytes out of all GFAP+ tanycytes with LiPR rescuing HFD-induced decrease at 7d (Bonferroni’s Test, Fig.2E). These data suggest that short LiPR administration in the context of HFD increases number and proportion of GFAP+ α-tanycytes. Because Vimentin is associated with nutrient transport in cells and with metabolic response to HFD ^52–54^, we quantified the proportion of GFAP+ tanycytes expressing Vimentin (Fig.2F). While treatment or duration of treatment had no effect on the absolute number of GFAP+Vimentin+ tanycytes (data not shown), the interaction between treatment type and duration of treatment had a statistically significant effect on the proportion of GFAP+Vimentin+ tanycytes (TWA, F(4,45) = 2.61, p = 0.048) with a statistically significant increase in the proportion of GFAP+Vimentin+ tanycytes at 7d (Bonferroni’s Test). In addition, a statistically significant increase in the proportion of GFAP+Vimentin+ tanycytes was observed in 21d Controls compared with 7d. Taken together, these results suggest that LiPR treatment in the context of acute HFD increases the density of GFAP+ α-tanycytes and the proportion of GFAP+Vimentin+ tanycytes. Because the increase in number of GFAP+ α-tanycytes could be due to increased cell proliferation, we stained brain sections for a marker of cell proliferation, Ki67 ^55,56^, and a marker of htNSCs, the Retina and anterior neural fold homeobox transcription factor (Rax).^51,57^ Unfortunately, we observed proliferating Ki67+Rax+ tanycytes so rarely (Fig.2G) that it did not allow reliable quantification. Nevertheless, we did not observe a statistically different change in the area occupied by Rax+ tanycytes (Fig.2H).

**Figure 2.**
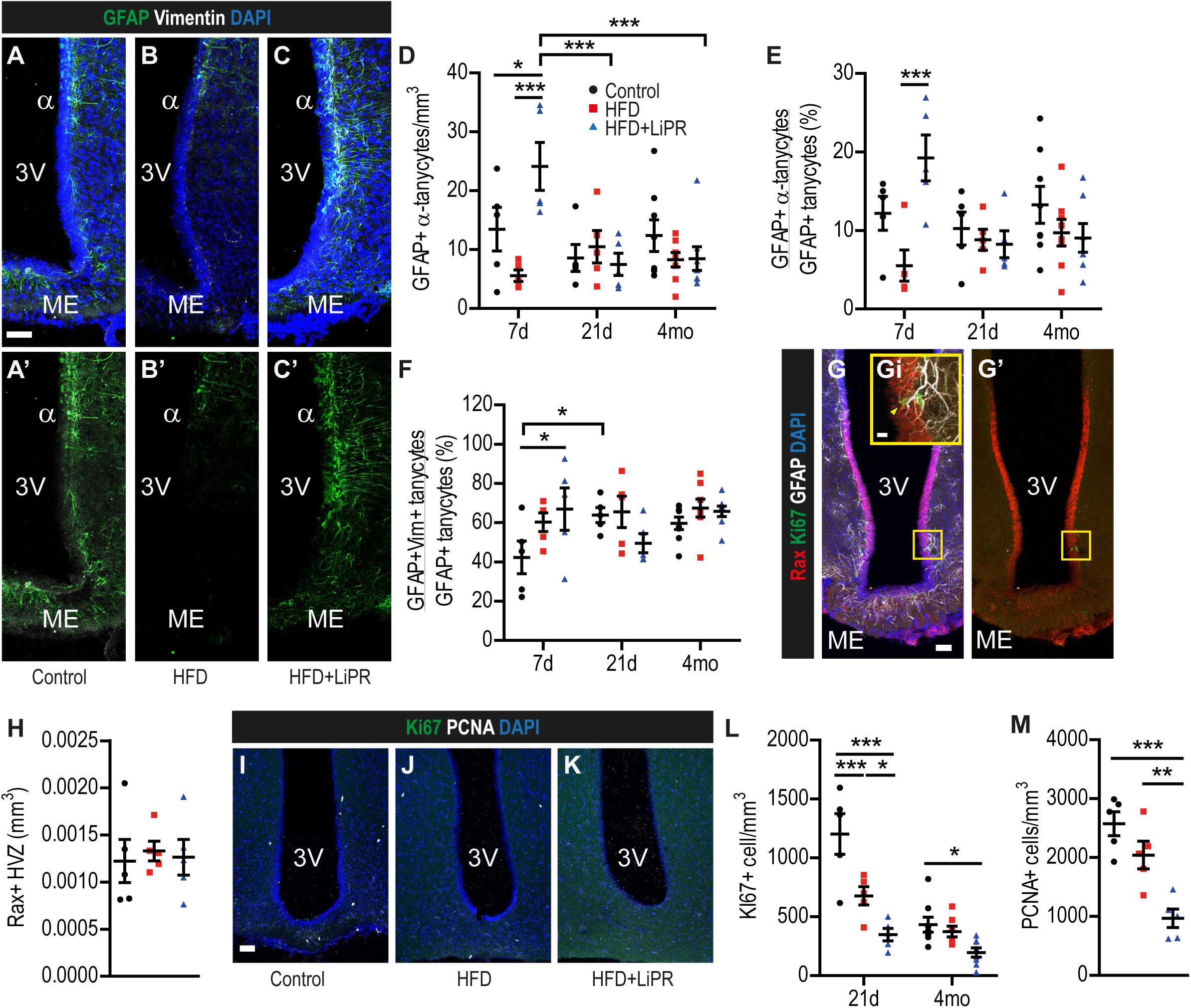
(A-C) Representative confocal images of HVZ with 3^rd^ Ventricle (3V) and Medial Eminence (ME) stained as indicated in Control (A), HFD (B) and HFD + LiPR (C) of 21d HFD group. (D) Quantification of GFAP+ α-tanycytes per volume of MBH. (E) Proportional analysis of GFAP+ α (F) Proportional analysis of GFAP+Vimentin+ tanycytes in MBH. (G) Representative image of HVZ stained as indicated. The insert (Gi) highlights a Rax+Ki67+GFAP+ tanycyte (yellow arrowhead). (H) Volume of Rax+ HVZ in the 21d HFD group. (I-K) Representative images of MBH stained as indicated (21d HFD group). (L-M) Quantification of MBH cells positive for Ki67 (K) and PCNA (L). Scale bars: 10 μm (Fi, M), 50 μm (all other). n = 5 for 7d and 21d groups, n = 8 for 4mo group. One-way ANOVA with Bonferroni’s test. *p < 0.05, **p < 0.01, ***p < 0.001. Data are presented as mean ± SEM.

Next, we administered Liraglutide to mice on HFD to determine if effects on tanycytes are a shared feature of anti-obesity treatment (Supplementary Figure 1A-C, S.Fig.1A-C). Liraglutide treatment in the 7d and 21d protocols did not change the number of GFAP+ α-tanycytes (S.Fig.1D) or proportion of GFAP+Vimentin+ tanycytes (S.Fig.1E) suggesting that unlike LiPR, Liraglutide does not affect number or character of tanycytes.

### Anti-obesity compounds reduce proliferation in the MBH

To determine if LiPR alters cell proliferation in the MBH (Fig.2I-K), we quantified cells expressing Ki67 and the Proliferating Cell Nuclear Antigen (PCNA).^58^ TW ANOVA revealed that treatment (F(2,32) = 24.8, p < 0.0001) and duration of treatment (F(1,32) = 40.81, p < 0.0001) had a statistically extremely significant effect on reduction of Ki67+ cells in the MBH (Fig.2L). In addition, an interaction between treatment type and duration of treatment had a statistically very significant effect on the number of Ki67+ cells (F(2,31) = 8.62, p = 0.001). Bonferroni’s Test showed that 21d (p < 0.001) but not 4mo HFD reduces number of Ki67+ cells and this reduction is magnified by LiPR administration in both protocols. In parallel, LiPR had a statistically very significant effect on reducing number of PCNA+ cells in 21d protocol (one-way ANOVA (OWA), F(2,12) = 16.66, p = 0.0003) when compared to both LFD and HFD groups (Fig.2M). These results suggest that LiPR decreases cell proliferation in the context of HFD. As with the analysis of tanycytes, we wanted to address if these effects on cell proliferation are shared among anti-obesity compounds. Indeed, administration of Liraglutide (S.Fig.2F-G) also very significantly decreased the number of both Ki67+ (TWA, F(2,26) = 20.96, p < 0.001) and PCNA+ cells (OWA, F(2,12) = 19.14, p < 0.0002) in the MBH, suggesting that different anti-obesity compounds exert similar effects on hypothalamic proliferation in the context of HFD.

### LiPR and Liraglutide display different effects on adult neurogenesis in the SGZ

Because obesity was found to alter adult neurogenesis in the SGZ ^59^, we determined effects of HFD, LiPR and Liraglutide in the hippocampal neurogenic niche (S.Fig.2A-D). Treatment with LiPR (TWA F(2,44) = 9.88, p = 0.0003), duration of treatment (F(2,44) = 6.86, p = 0.0002) and interaction of treatment and its duration (F(4,44) = 9.88, p = 0.0003) had statistically extremely significant effects on reduction of Ki67+ cells in the SGZ, with the 21d HFD showing statistically significant reduction in proliferation, which was not rescued by LiPR (S.Fig.2E). Similarly, number of PCNA+ cells in the SGZ was extremely affected by LiPR treatment (TWA F(2,45) = 21.19, p < 0.0001), duration of treatment (F(2,45) = 374.3,) and their interaction (F(4,45) = 9.88, p = p < 0.0001) with HFD-induced reduction in PCNA+ cells in the 21d protocol, which was further reduced by LiPR administration (S.Fig.2F). In contrast to decreasing proliferation, LiPR rescued HFD-induced reduction in number of DCX+ neuroblasts (S.Fig.2G) and DCX+ neurons (S.Fig.2H). For DCX+ neuroblasts, there was a statistically extremely significant effects of LiPR treatment (TWA F(2,45) = 11.38, p < 0.0001), duration of treatment (F(2,45) = 463.03, p < 0.0001) and their interaction (F(4,45) = 6.25, p = 0.0004) with Bonferroni’s Test revealing HFD-induced reduction in the number of DCX+ neuroblasts, which was rescued in the 21d protocol. Similarly, DCX+ neurons in the SGZ were statistically significantly affected by treatment (F(2,45) = 8.77, p = 0.0006), duration of treatment (F(2,54) = 552.73, p < 0.0001) and interaction of treatment and its duration (F(4,45) = 3.78, p = 0.011). There was a statistically significant HFD-induced decrease in the number DCX+ neurons, which was rescued by LiPR administration in the 21d protocol (Bonferroni’s Test). Analysis of Liraglutide-treated mice revealed difference between these two anti-obesity compounds. In contrast to LiPR, Liraglutide rescued HFD-induced reduction in Ki67+ cells in 21d (S.Fig.2I), however, failed to rescue the HFD-induced decrease in number of DCX+ neuroblasts (S.Fig.2J) and even further reduces the number of DCX+ neurons (S.Fig.2K). TW ANOVA showed that treatment (F(2,24) = 4.3, p = 0.025) and duration of treatment (F(1,24) = 4.22, p = 0.05) but not their interaction have statistically significant effects on number of Ki67+ cells. Similarly, treatment (F(2,24) = 7.21, p = 0.0035) and duration of treatment (F(1,24) = 95.86, p < 0.0001) but not their interaction have statistically significant effect on DCX+ neuroblasts or DCX+ neurons (treatment (F(2,24) = 12.29, p = 0.0002, duration of treatment F(1,24) = 161.57, p < 0.0001, and their interaction F(2,24) = 5.96, p = 0.0079). These results suggest that LiPR reduces proliferation but increases neuronal differentiation in the SGZ, whereas Liraglutide has the opposite effects. This also suggests that LiPR can rescue neuronal differentiation reduced by the HFD in the SGZ and that, in both neurogenic niches, it reduces cell proliferation.

### LiPR treatment generates smaller MBH-derived neurospheres

The pool of proliferating cells in the MBH consists of not only proliferating htNSCs but also neural and glial precursors.^60^ To observe the effects of LiPR on proliferation and cell cycle properties of htNSC, we treated mice with Control Diet or HFD with or without concurrent administration of LiPR for 14 days. After the treatment, primary htNSCs were grown as globular syncytia, so-called neurospheres ^35,61^, or as adherent cell cultures followed by time-lapse imaging.^61^ MBH-derived neurospheres (Fig.3A-C) from HFD-treated mice showed no difference in their diameter after 5d (Fig.3D,E) or 10d (Fig.3F,G) when compared with Controls. However, neurospheres from animals exposed to HFD+LiPR had smaller diameter and showed size distribution skewed towards smaller neurospheres at both 5d (OWA, F(2,73) = 15.80, p < 0.0001) and 10d (Kruskal-Wallis test, H = 10.30, p = 0.0058). While the LiPR-treated neurospheres were smaller, there was no difference in the number of neurospheres in any of the treatment groups (S.Fig.3A-B). Interestingly, neurospheres had smaller diameter also from animals treated with LiPR but kept on Control diet (S.Fig.3C-D) but only in 5d and not 10 days in culture (S.Fig.3E-F). *In vivo* treatment with LiPR followed by 10 days in culture resulted in higher qPCR expression of Prlh, a gene that encodes PrRP, in context of HFD (Fig.3H) and Control diet (S.Fig.3G) but not in changes of expression of Prlhr, encoding GPR10 receptor (Fig.3H) or Vimentin (S.Fig.3I). These results suggest that LiPR reduces cell proliferation both *in vivo* and *in vitro* and increases PrRP expression in MBH-derived stem cells and progenitors.

**Figure 3.**
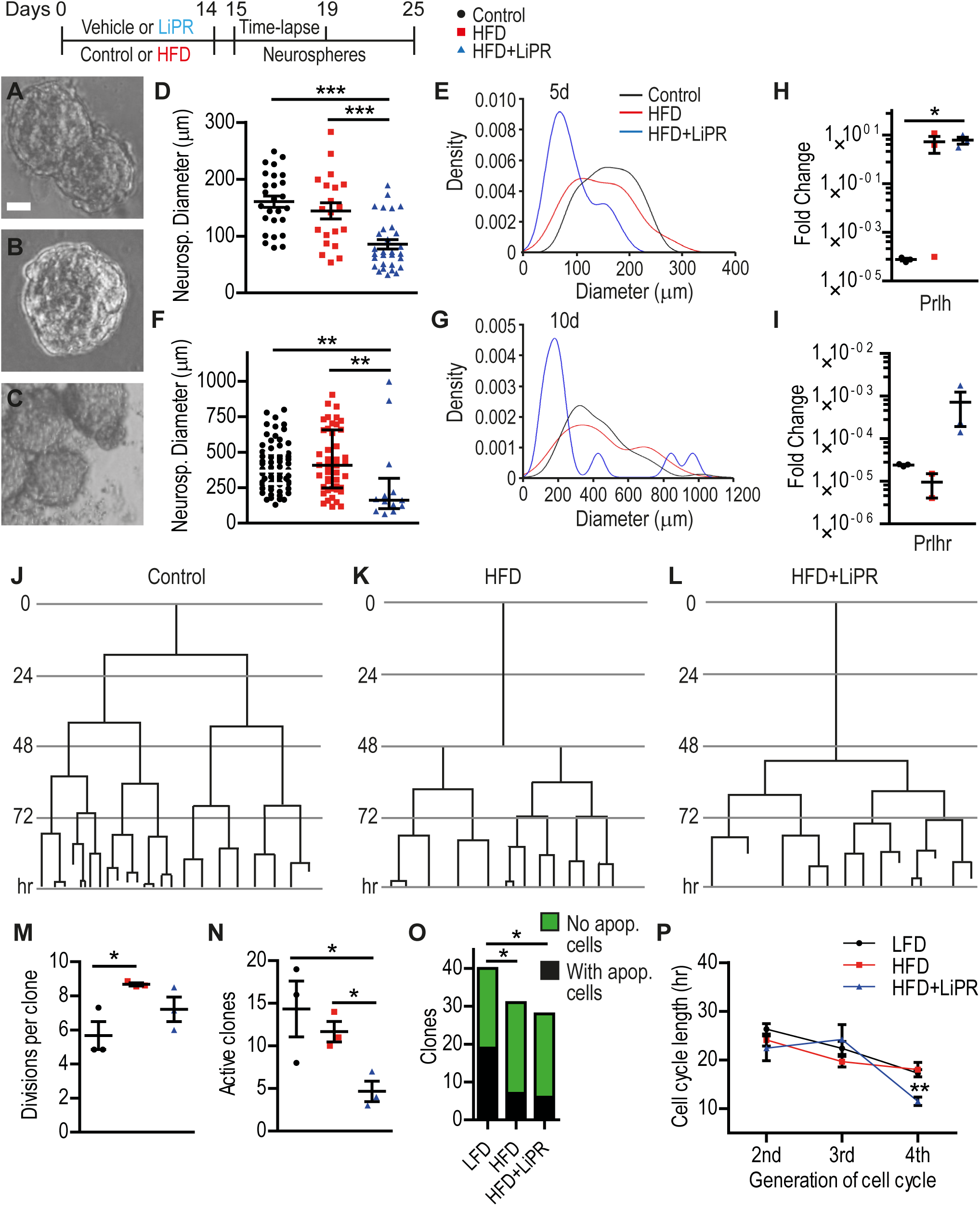
(A-C) Representative images of HVZ-derived neurospheres 5d in culture from Control (A), HFD (B) and HFD + LiPR (C) treated mice. Schematics of the experimental protocol shown above. (D,F) Quantification of diameter of neurospheres 5d (D) and 10d (F) in culture. (E,G) Kernel density plots of neurosphere diameter frequency distribution as a function of diameter for 5d (E) and 10d (G) in culture. (H-I) Relative mRNA fold change of Prlh (H) and Prlhr (I) compared to Gapdh in cDNA from MBH-derived neurospheres (10d in culture). (J-L) Example cell division trees from 4-day time-lapse imaging of aNSCs from HVZ of Control (J), HFD (K), HFD + LiPR (L) mice. (M) Quantification of the number of cell divisions per division tree clone. (N) Time-lapse quantification of active (dividing) clones per 20.000 plated cells. (O) Proportion of active clones containing at least one apoptotic cell. (P) Cell cycle length from the time-lapse imaging over observed cell divisions. Scale bars: 20 μm. n = 3. Chi-square test (O), Kruskal-Wallis test with Dunn’s test (F), or one-way ANOVA (all other) with Bonferroni’s test. *p < 0.05, **p < 0.01, ***p < 0.001. Data are presented as median ± IQR (F) or mean ± SEM (all other).

### LiPR affects cell activation and cell cycle of htNSCs

To further understand effects of LiPR on cell cycle and proliferation, we performed time-lapse imaging *in vitro*. Individual htNSCs were continuously imaged to obtain cell division trees from the *post-hoc* analysis (Fig.3J-L). In total, we traced 48 individual cells from Control Diet, 68 from HFD and 53 from HFD + LiPR groups. The analysis revealed that treatment significantly increase number of cell divisions per clone (OWA, F(2,6) = 5.61, p = 0.042), which is significant for HFD but not HFD+LiPR (Fig.3M). In addition, LiPR treatment reduced number active clones (OWA, F(2,6) = 5.47, p = 0.044) defined as clones with at least one cell division during the time-lapse imaging window (Fig.3N). As with the neurospheres, we observed effects of LiPR also in the context of Control diet (S.Fig.3J; 25 individual cells traced), where it does not change number of cell divisions (S.Fig.3K) but decrease number of active clones (S.Fig.3L), an effect clearly independent of diet (S.Fig.3M). However, this diet independent effect on cell activation is not mirrored by LiPR’s effect on cell death and cell cycle. Time-lapse imaging reveals that there are fewer clones with at least one apoptotic cell in both HFD and HFD+LiPR (Fig.3O) but not Control diet + LiPR (S.Fig.3N). Furthermore, the generation of observed cell cycle from 1^st^ to 4^th^ a statistically a very significant effect on cell cycle length (TWA, F(2,237) = 16.02, p < 0.0001), eventually resulting in shorter cell cycle in HFD + LiPR cells (Fig.3P), which is in contrast with extension of cell cycle in clones from LiPR treated mice on Control diet (S.Fig.3O). Of note, our data for the first time directly show the length of cell cycle in htNSCs (22 ± 0.78 hrs for 2^nd^-4^th^ cell divisions), which is significantly shorter than cell cycle of aNSCs from the adult Subventricular Zone, SVZ (28.55 ± 2.14 hrs; unpaired, two-tailed T-Test, p = 0.0025) ^61^ but longer than cycle length of aNSCs from the SGZ (16.62 ± 3.28 hrs; unpaired, two-tailed T-Test, p = 0.021) ^62^ observed by *in vitro* time-lapse imaging under the same proliferating conditions. Finally, we tested whether PrRP signalling is cell intrinsic to htNSCs. Because LiPR is highly adhesive to cell culture plasticware, we added recombinant human PrRP31 (hPrRP31) protein, which is not plastic adhesive, to WT primary htNSC and determine its effects on cell dynamics and proliferation. Exposure to hPrRP31 *in vitro* significantly reduced number of cells per clone and cell divisions (S.Fig.4A-D) suggesting that PrRP signalling acts directly on htNSCs to reduce their proliferation. Taken together, these results suggest that LiPR and PrRP decrease activation and proliferation of htNSCs and changes their cell cycle in diet-dependent manner.

### LiPR rescues HFD-induced decrease in survival of new MBH neurons

Our next goal was to determine whether LiPR or Liraglutide change survival of adult-generated cells in the MBH. We exposed mice with three different BrdU protocol with varying length of post-BrdU chase periods. In the 7d and 21d HFD protocols, there was 6d and 16d chase periods, respectively (S.Fig.5). In the 7d protocol, there was no difference between any of the treatment groups (Control, HFD, HFD+LiPR and HFD+Liraglutide) in number of BrdU+ cells, neurons, or astrocytes (S.Fig.5A-G). In the 21d protocol (S.Fig.5H-K), LiPR did not change the number of cells, neurons, or astrocytes positive for BrdU in the context of HFD (S.Fig.5L-N). However, in the Arcuate Nucleus (ArcN), the primary nutrient and hormone sensing neuronal nucleus of MBH ^4^, there was a statistically significant increase of BrdU+ neurons by HFD compared to Control (data not shown) and Liraglutide, in context of HFD, decreased the number of BrdU+ cells compared to Control (S.Fig.5L; Kruskal-Wallis test, H = 8.71, p = 0.033, Dunn’s post-test, p < 0.05) and decreased the number of BrdU+ astrocytes compared to HFD (S.Fig.5N; OWA, F(3,16) = 4.3, p = 0.0021, Bonferroni post-test, p < 0.05). Interestingly, LiPR, in context of Control diet, significantly decreased number of BrdU+ cells (S.Fig.5O; p = 0.0015) without altering the number of BrdU+ neurons or astrocytes (S.Fig.5P-Q). Taken together, these results suggest that Liraglutide in the context of HFD and LiPR in the context of Control diet reduces the number of new BrdU+ cells in the MHB but only in the 21d and not 7d protocol. These results also indicate that Liraglutide but not LiPR decrease survival of new cells in the MBH in the context of week to 3-week exposure to HFD.

**Figure 4.**
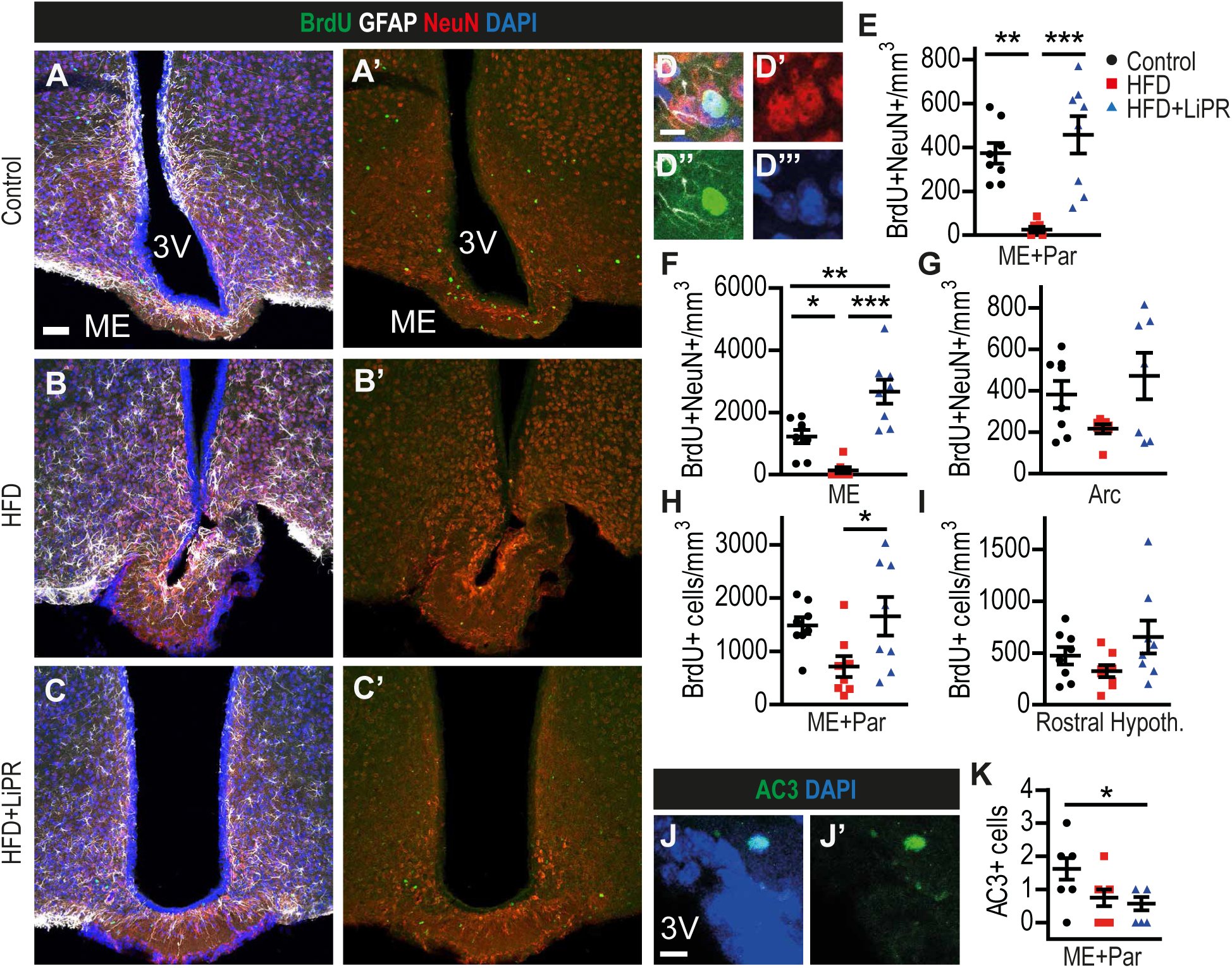
(A-C) Representative confocal images of HVZ stained as indicated in Control (A), HFD (B) and HFD + LiPR (C) of 4mo HFD group. (D) An example of BrdU+ neuron in the MBH parenchyma. (E-G) Quantification of BrdU+ neurons in MBH parenchyma (E), ME (F) and Arcuate Nucleus (G). (H-I) Quantification of all BrdU+ cells in parenchyma of MBH (H) and rostral hypothalamus (I). (J) An example of Activated Caspase 3 (AC3) positive cell near 3V wall. (K) Quantification of AC3+ cells in MBH. Scale bars: 50 μm (A-C), 10 μm (D). n as in Fig.1. One-way ANOVA with Bonferroni’s test. *p < 0.05, **p < 0.01, ***p < 0.001. Data are presented as mean ± SEM.

**Figure 5.**
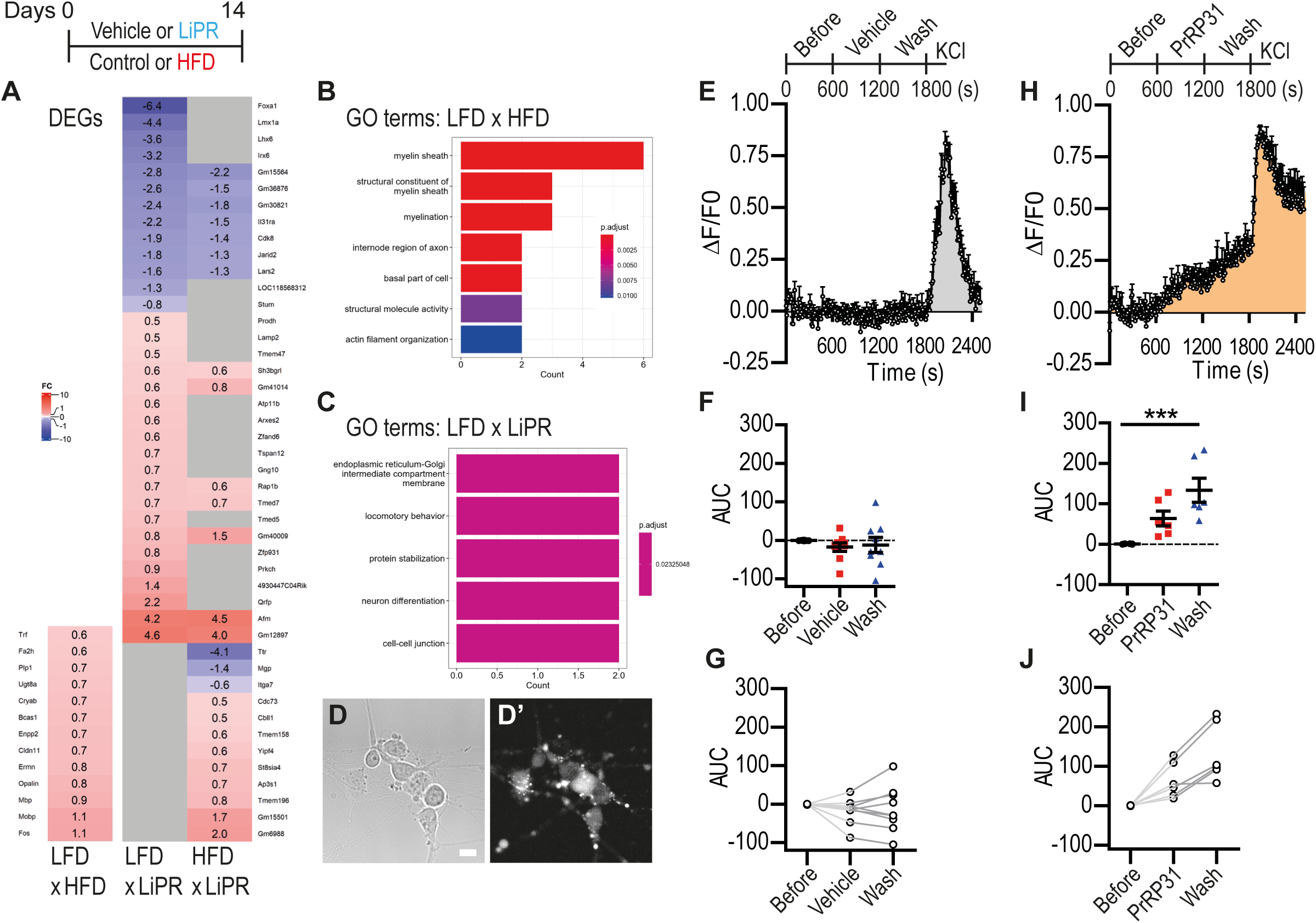
(A) A table of Differentially Expressed Genes (DEGs) in 3 different pair-wise comparisons of bulk RNAseq data. Experimental protocol is depicted above. (B) Bulk RNAseq Gene Ontology (GO) terms for the LFD vs HFD comparison. (C) Bulk RNAseq GO terms for LFD vs LFD+LiPR comparison. (D) A representative image in the bright field (D) and fluorescence (D’) of hiPSC-derived hypothalamic neurons loaded with the Rhod-3 AM dye and used in calcium imaging. (E-H) A graph of the relative fluorescence change (ΔF/F0) of Rhod-3 as a function of time before and during hPrRP31 (or Vehicle) application in medium, wash-out and KCl positive control. (E) Neurons exposed to Vehicle. (H) Neurons responding to hPrRP31. (F-I) A summary of fluorescence before (0-600 s), during hPrRP31 (F) or Vehicle (I, 600- 1200 s) or wash-out (1200-1800 s) in Area Under Curve (AUC). (G-J) Individual neurons from calcium imaging in the before-after plot from the Vehicle (G) and hPrRP31-responsive (J) groups. Scale bar: 10 μm. n = 3-4 for RNAseq, n = 9 for Control and n = 6 for hPrRP31-responsive neurons. One-way ANOVA with Bonferroni’s test, ***p < 0.001 (F,I). Data are presented as mean ± SEM.

While short-term exposure to HFD can alter functions of the MBH ^46^, it does not result in DIO (Fig.1A-C). To test the effects of DIO on survival of newly generated cells in the MBH, mice exposed to 4mo HFD were administered BrdU for 2wk (on day 92) followed by 2wk chase (Fig.1C). Exposure to HFD dramatically reduced the number of new BrdU+ neurons, which was rescued by LiPR administration (Fig.4A-D). OWA revealed an extremely significant effect of the treatment on number of BrdU+ neurons with a statistically very significant decrease in the number of BrdU+ neurons by HFD (Bonferroni’s Test, p < 0.01) rescued by LiPR (p < 0.001) to the level of Controls (Fig.4E). This statistically significant rescue of HFD-induced decrease in number of BrdU+ neurons by LiPR was observed not only in MBH parenchyma (Fig.4E; OWA, F(2,20) = 14.21, p < 0.0001) but also in the ME (Fig.4F; OWA, F(2,20) = 21.37, p < 0.0001) but not in ArcN (Fig.4G). This rescue effects is not limited only to new neurons but also to other BrdU+ cells (Fig.4H; OWA, F(2,21) = 3.87, p = 0.037) and it is confined to the MBH as there is no difference in the number of BrdU+ cells between Control, HFD or HFD+LiPR in the rostral hypothalamus (Fig.4I). These results suggest that LiPR promotes survival of new MBH neurons in the context of DIO, which could be caused by reduced cell death. To test this alternative, we quantified the number of cells in the 3V walls and MBH parenchyma positive for the Activated Caspase 3 (AC3), a marker of apoptosis.^63^ Indeed, our results show (Fig.4J,K) a statistically significant decrease in number of AC3+ cells (OWA, F(2, 22) = 2.46, p =0.025) in HFD+LiPR compared to HFD (Bonferroni’s Test, p < 0.05).

### LiPR displays no effects on astrocytes

Because obesity increases reactivity of glial cells ^46,64^, we determined the effects of HFD and LiPR on astrocytes. In the 4mo HFD protocol, OWA revealed a statistically extremely significant effects of HFD exposure on number of GFAP+ astrocytes in the MBH (F(2,21) = 16.22, p < 0.0001), which was not altered by LiPR (S.Fig.6A-D). Similarly, HFD increases the optical density of GFAP in the MBH independently of LiPR (S.Fig.6E; OWA, F(2,21) = 5.67, p = 0.011). However, this HFD-induced increase in reactive astrogliosis is not mirrored by an increase in astrogliogenesis as there are no effects of HFD or LiPR on number of new BrdU+ astrocytes in the MBH (S.Fig.6F,G), suggesting that LiPR does not alter either reactive astrogliosis or astrogliogenesis and its pro-survival effects are confined to new neurons (Fig.4).

### LiPR administration alters gene expression in the context of diet

To elucidate molecular mechanisms of action of LiPR, we performed bulk RNA sequencing (RNAseq) of the MBH. This brain region was chosen because it contains the HVZ and the hypothalamic parenchyma, where we observed the changes in number of htNSCs and proliferating and surviving adult-generated cells. In addition, the MBH has a high density of cells expressing PrRP receptors (Fig.1L-Q). The MBH tissue was sampled from mice exposed to Control or HFD with concurrent LiPR administration for 14 days (Fig.5). Analysis of Differentially Expressed Genes (DEGs, Fig.5A) revealed that LiPR administration changes expression of genes involved in cell cycle regulation (e.g. Cdk8, Cdc73, Jarid2) or in function of the Golgi apparatus (e.g. Ap31s, Tmed7, Yip4), supporting the time-lapse results that suggest LiPR influences cell cycle and activation of htNSCs. In addition, LiPR changed expression of individual DEGs with unique functions in insulin expression (Lmx1a), food intake (Qrfp), tissue inflammation (Il31ra) or thyroid hormone and vitamin dynamics (e.g. Afm, Ttr). These genes are involved in controlling metabolism and energy homeostasis. Gene Ontology (GO) analysis of genes with changed expression (Fig.5B-C, S.Fig.7A-B) highlighted GO terms involved in neuron differentiation, cell-cell junction, or endoplasmic reticulum-Golgi compartments, suggesting that LiPR may be involved in these processes. Finally, comparison of Control diet with HFD without LiPR administration showed GO terms strongly involved in myelination, myelin sheath and its constituents, actin filament organization and internode region of axons. Several upregulated DEGs from this comparison were involved in myelination (e.g. Mbp, Mobp, Cldn11, Ugt8a, Plp1). This suggests that HFD promotes myelination, which is, however, not affected by LiPR as there are no myelin related DEGs in the pair comparison between Control diet or HFD and LiPR groups. RT-qPCR from bulk RNA of MBH confirmed expression changes of selected genes observed by RNAseq (S.Fig.7C) and showed that HFD increases expression of Prlh, which encodes the native PrRP protein, but not Prlhr, encoding GPR10 (S.Fig.7D).

### PrRP increases intracellular calcium in human hypothalamic neurons

Our final goal was to determine whether PrRP can influence physiology in neurons. This should address physiological responsiveness of the target cell population. As a model, we chose human induced Pluripotent Stem Cell (hiPSC)-derived hypothalamic neurons ^65,66^ and determined their responsiveness to PrRP analogues by calcium imaging. Differentiated hiPSC-derived hypothalamic neurons were loaded with Rhod-3 AM dye (Fig. 5D) and calcium-induced changes in fluorescence in individual neurons were recorded over time before and during the application of hPrRP31, which was used instead of LiPR because it is not plastic adhesive (see also S.Fig.4). In contrast to vehicle-treated control neurons (Fig. 5E-G), approximately half of neurons exposed to hPrRP31 (46 %, 6 out of 13) displayed a robust and steady increase in intracellular calcium (Fig. 5F). One-way ANOVA showed a statistically significant increase in calcium-dependent fluorescence (F(2,15) = 10.90, p = 0.0012) during hPrRP31 application (unpaired, two-tailed T-Test, p = 0.038) that was retained after wash-out (Bonferroni’s Test, p < 0.001; Fig. 5G-H). This hPrRP31 effect on intracellular calcium was not present in non-responsive neurons (S.Fig.8A-C), which diminished the effects when pooled together with the hPrRP31-responsive neurons (S.Fig. 8D-F). These results demonstrate that PrRP signalling stimulates increase in the intracellular calcium in hiPSC-derived hypothalamic neurons suggesting their direct responsiveness to PrRP and LiPR.

## Discussion

### LiPR decreases body weight and promotes expression of PrRP receptors

For the first time, our results show that anti-obesity compounds influence adult neurogenesis. We demonstrate that administration of a PrRP analogue increases number of htNSCs, decreases cell proliferation and improves survival of new neurons. Interestingly, our results indicate that different anti-obesity compounds do not exert same effects on adult neurogenesis. In agreement with previous results ^15,16^, our data demonstrate that LiPR exerts anorexigenic effects by reducing the body weight and decreasing plasma concentrations of insulin, leptin, and cholesterol that were increased by the HFD. This ability of LiPR to normalize metabolomic response to HFD is further highlighted by our RNAseq data. LiPR upregulated expression of genes related to energy homeostasis and development of obesity such as Qrfp and Afm ^67,68^ or genes involved in regulation of insulin or thyroid hormone expression and distribution such as Lmx1a and Ttr.^69,70^ In addition to its role in metabolism, LiPR also downregulated expression of pro-inflammatory cytokine, Il31ra ^71^ and influenced expression of Tmed7, which controls trafficking of Toll-like receptor 4 (Trl4), a key receptor in innate immunity.^72^ This suggests that LiPR plays an anti-inflammatory role previously observed in a model of neurodegeneration.^73^ Our results also improve the understanding of expression of PrRP receptors in the hypothalamus, which has been demonstrated previously but without the cellular resolution or without co-localization with specific cellular markers.^48,74–76^ Interestingly, our data show that LiPR can dynamically regulate PrRP signalling by rescuing HFD-induced decrease in GPR10 protein density in the MBH and by increasing PrRP expression both in the MBH and in MHB-derived neurospheres. This LiPR-induced increase in the expression may suggest a positive feedback of the agonist on its signalling pathway as it is the case with estrogen-induced estrogen secretion.^77^ Or it may indicate a desensitisation of GRP10 signalling by LiPR, which would resemble insulin resistance in T2DM.^78^ Importantly, we show for the first time that GPR10 and NPFF2 receptors are expressed in the MBH-derived neurospheres and localize in new adult-generated hypothalamic neurons suggesting that PrRP signalling can act directly on adult neurogenesis, which is what we investigated here.

### LiPR increases number of htNSCs by reducing their activation

Our results from cell quantification in the MBH and the SGZ and primary MBH-derived cell cultures jointly suggest that LiPR promotes adult neurogenesis. In the context of HFD, we show that LiPR increases number and proportion of GFAP+ α-tanycytes, which are considered the putative htNSCs.^36,38,79^ Increasing the number of GFAP+ α-tanycytes cannot be due to their higher proportion because the absolute number of GFAP+ tanycytes or the area of HVZ covered by Rax, which does not cluster with GFAP+ in tanycytes ^80^, do not change. This increase could be explained by lower cell death or higher self-renewal proliferation that would increase the maintenance and population of htNSCs.^28^ However, the time-lapse imaging showed no reduction in number of clones containing apoptotic cells from HFD+LiPR vs HFD groups and HFD+LiPR decreased rather than increased cell proliferation both *in vivo* and *in vitro*. This suggests mechanisms such as reduced cell activation. Indeed, our time-lapse imaging results show that LiPR decreases number of active clones and shortens cell cycle length, which could be a compensatory mechanism for the lower activation. The lower cell activation may protect the stem cell pool from exhaustion elicited by the HFD. In addition, our observation that adult NSCs from different niches have different duration of the cell cycle *in vitro* indicates that it is intrinsically controlled. The role of LiPR in regulating cell cycle is also supported by our findings that it reduces proliferation both in the MBH and the SGZ and influences the expression of genes that control the cell cycle (e.g. Cdk8, Jarid2).^81,82^ Interestingly, recent RNAseq data from mouse macrophages exposed to PrRP for only 18 hours reveal DEGs related not only to neuronal differentiation but also to cell cycle or inflammation, such as Cdkn2a or Il1b.^83^ In addition, we show that HFD+LiPR also increases the proportion of Vimentin+GFAP+ tanycytes, a subset of α-tanycytes, which may be protective against HFD-induced changes as Vimentin is responsive to HFD ^52^ and is downregulated in tanycytes of mice with a hypermetabolic phenotype.^54^ Taken together, our results improve our understanding of how the HFD and the anti-obesity pharmacology influence tanycytes, which act not only as htNSCs but as metabolic regulators that control vascular permeability, leptin transport ^85^ and the entry of anti-obesity compounds such as Liraglutide to the hypothalamic parenchyma.^37^

### LiPR rescues reduced survival of new MBH neurons

The neurogenic process requires not only active adult NSCs but also survival of new adult-generated neurons. The exposure to the HFD was shown to increase or decrease hypothalamic adult neurogenesis depending on the length of HFD treatment and timing of cell birth-dating protocols.^31,32,34,35,42^ Our results show that the short and intermediate exposure to HFD does not change the number of newly generated, BrdU+ cells, neurons, or astrocytes in the MBH. These findings are in agreement with previous studies that used short and intermediate HFD protocols ^31,42^, which can induce reactive gliosis or inflammation but not obesity.^46,86^ In addition, our results show that LiPR does not change the number of new cells, neurons, or astrocytes, suggesting that it does not alter cell survival in the context of short HFD exposure. This is in contrast with the long HFD exposure that induces obesity and reduces survival of new hypothalamic neurons both in our results and previous studies.^34,35^ Importantly, we show that LiPR rescues the reduction in the number of new neurons caused by the long HFD exposure. This suggests that LiPR increases their survival. These pro-survival effects, however, are confined to the MBH as there is no difference in the number of new cells in the rostral hypothalamus. Because DIO can increase cell death in the adult hypothalamus ^34,87^, we also determined if LiPR can reduce apoptosis. Indeed, our results show that LiPR reduces the number of AC3+ cells in the MBH suggesting that the increased number of new neurons is thanks to lower apoptosis. Alternatively, LiPR could rescue neuronal survival also by mitigating HFD-induced reactive gliosis, which contributes to the increased inflammation and cell death.^46,88^ However, we did not observe that LiPR mitigated gliosis suggesting that the improved neuronal survival is a cell-intrinsic feature.

### LiPR activates human hypothalamic neurons

To determine whether LiPR can directly activate new neurons, we performed calcium imaging. PrRP or LiPR were shown to increase intracellular calcium in cell lines expressing human GPR10 ^27,48,89^ and to reduce the frequency of GABAergic miniature Inhibitory Postsynaptic Currents (mIPSCs) in murine hypothalamic neurons.^90^ This suggests that PrRP signalling decreases neuronal inhibition. However, to our best knowledge, physiological responsiveness to PrRP has not been demonstrated in immature or human neurons.^20^ To fill this knowledge gap, we performed calcium imaging in hiPSC-derived neurons. We chose these neurons for two principal reasons. First, they display immature characteristics when compared to the tissue-derived counterparts ^91–93^, which makes them relevant as a model of immature, adult neurogenesis-generated neurons. Second, they provide relevance to human biology and medicine. Specifically, hiPSC-derived hypothalamic neurons display MBH lineage and phenotype ^65,66^ and thus are ideal for testing responsiveness to PrRP. This is especially relevant considering that highest concentration of PrRP in the human brain was detected in the hypothalamus.^94^ Our results from calcium imaging show for the first time that PrRP can directly stimulate human iPSC-derived hypothalamic neurons. Interestingly, the calcium concentration increases even when hPrRP31 was washed-out. This might be caused by persistent signalling downstream of PrRP receptors or by internalization of the receptors. Finally, almost half of hypothalamic neurons randomly chosen for calcium imaging displayed responsiveness to PrRP. However, anorexigenic POMC-expressing neurons, which are expected to be primary targets of PrRP, represent less than 10% of total hypothalamic neurons differentiated from hiPSCs.^65^ This suggest that other hypothalamic neuronal subtypes may be responsive to PrRP as well.

### LiPR rescues HFD-induced decrease in SGZ neurogenesis

Because diet influences adult neurogenesis in the hippocampus ^95^, we determined the effects of HFD and LiPR in the SGZ. In the context of the intermediate HFD protocol, we observed that LiPR reduces cell proliferation already lowered by the HFD, whereas it rescues reduction in number of neuroblasts and new immature neurons, which was induced by the HFD, a finding reported previously in the SGZ.^96,97^ These results suggest that LiPR exhibits pro-neurogenic functions in both the HVZ and the SGZ, which is in line with its other neuroprotective effects.^98^ The dual effects of LiPR on two neurogenic niches are especially intriguing given the fact that the hypothalamus can modulate adult neurogenesis in the hippocampus.^99^

### Diet independent effects of LiPR on hypothalamic neurogenesis

As LiPR displays a broad range of roles independent from the HFD ^22,100^, we determined its effects on hypothalamic neurogenesis in the context of Control diet. In the MBH, LiPR dramatically reduced number of new, adult-generated cells, which is not a consequence of lower cell survival, as the Control diet is not expected to induce cell death, but rather a consequence of reduced cell proliferation. Indeed, we observed that LiPR decreases proliferation also *in vitro* together with increasing the expression of Prlh and decreasing the activation of the htNSCs. These results jointly suggest that certain LiPR effects on htNSCs are diet independent. Importantly, *in vitro* activation of PrRP signaling in naïve WT htNSCs reduces cell proliferation, which supports the conclusion that LiPR actions on htNSCs are cell intrinsic.

### Liraglutide does not share neurogenic effects with LiPR

Given the effects of LiPR on adult neurogenesis, we were intrigued if anti-obesity compounds share the neurogenic potential as many anti-depressants do.^43,44^ Contrary to our expectation, our results show major differences between LiPR and Liraglutide, a compound already approved for human medicine, which is indicated for T2DM and has anti-obesity activity.^11,101^ While both compounds reduce cell proliferation in the MBH, Liraglutide, unlike LiPR, does not increase the number of htNSCs. Importantly, Liraglutide does not rescue HFD-driven reduction in the number of neuroblasts and immature neurons in the SGZ. Instead, it further decreases their number in the context of the HFD. These results point towards a conclusion that Liraglutide, a GLP-1RA receptor agonist, does not exert the same neurogenic effects as LiPR, an agonist of GPR10 and NPFF receptors. The difference in their action may originate in the difference of the cellular expression profile and signaling in their respective pathways. Nevertheless, this dissimilarity in neurogenic effects is intriguing because both compounds share other functions such as neuroprotection.^73^ More importantly, the positive effects of LiPR on adult neurogenesis warrant further research because healthy neurogenesis is essential for many brain functions such as learning and mood control.^44^

## Methods

### Mice

Animal experiments in this study were performed either in the United Kingdom (UK) or in the Czech Republic as it is specified below. All experimental procedures in the in UK were performed in accordance with the United Kingdom’s the Animals (Scientific Procedures) Act 1986 (ASPA) and were approved by the UK Home Office (HO) and the Biological Services of the College of Biomedical and Life Sciences, Cardiff University. All mice were 8-9 weeks old at the start of the regulated procedures. All mice were C57BL/6j males purchased from a HO accredited vendor, Charles River, UK. After transfer to our animal facility, mice were acclimatized for a minimum of 5 days before entering a regulated procedure. Mice were housed at a constant 22 °C, a relative humidity of 45 – 65 % and in a 12:12 hour light-dark cycle and group housed (3-5 mice per group) with *ad libitum* access to water and diet. Mice used for the *in vitro* experiments, time-lapse imaging and RNAseq were treated in the UK.

In the Czech Republic, all animal experiments followed the ethical guidelines for animal experiments and the Act of the Czech Republic Nr. 246/1992 and were approved by the Committee for Experiments with Laboratory Animals of the Academy of Sciences of the Czech Republic. C57BL/6 male mice that were 4 weeks old were obtained from Charles River Laboratories (Sulzfeld, Germany). The mice were housed under controlled conditions at a constant temperature of 22 ± 2°C, a relative humidity of 45 – 65% and a fixed daylight cycle (6 am – 6 pm), with 5 mice per cage. The animals were provided free access to water and the standard rodent chow diet. The 7d, 21d and 4mo protocols with BrdU treatment were conducted in the Czech Republic.

### Diets

Mice were given either Control Diet (Low Fat Diet, LFD) or High Fat Diet (HFD) in form of hard pellets. HFD was custom-made in the laboratory based on ^15^ and had energy content of 5.3 kcal/g, with 13%, 60% and 27% of calories derived from proteins, fats, and carbohydrates, respectively. This HFD is based on Ssniff Germany Diet (Ssniff^®^ R/M-H #V1530-000 RM-Haltung, Mehl (ground/powder), Ssniff Spezialdiäten GmbH, Soest, Germany), which is the basis of the Control (LFD) pellet diet (#V1535 RM-Haltung, 15 mm pellets, with 24%, 7%, 67% of calories derived from proteins, fat, and carbohydrates, respectively). HFD composition was 400 g of Ssniff base, 340 g of powdered cow-milk-based human baby formula (Sunar Complex 3, Dr Max Pharmacy, Mirakl, a.s., Bratislava, Slovakia), 10 g of Corn Starch and 250 g of pork lard (Lidl UK, Surbiton, UK). Ingredients were thoroughly mixed by a table-top kitchen robot, made into pellets and stored at −80 °C. Before there were given to animals, pellets were warmed to room temperature. Both control and HFD pellets were changed twice per week.

### Prolactin Releasing Peptide and its lipidized analogue

Full-length, 31 amino acid long human Prolactin Releasing Peptide (hPrRP31), SRTHRHSMEIRTPDINPAWYASRGIRPVGRF-NH_2_ was used in selected *in vitro* experiments as described below. Purified, desicated, crystal hPrRP31 (M.W. = 3665.15) was stored at −20 °C. Stock solution of 1 mM hPrRP31 in water was added to cell media to the final concentration of 1 μM. A human palmitoylated analogue of PrRP (LiPR) with the sequence SRTHRHSMEIK (N-γ-E(N-palmitoyl)) TPDINPAWYASRGIRPVGRF-NH_2_ was synthesized and purified at the Institute of Organic Chemistry and Biochemistry, Czech Academy of Sciences, Prague (CAS), Czech Republic, as previously described ^16^. Purified, desicated, crystal LiPR (M.W. = 4004) was stored at −20 °C until it was reconstituted in the sterile physiological solution (0.9% NaCl, pH = 7.4) as 1 mg/ml concentration (corresponding to approx. 0.25 mM). This working stock solution was aliquoted to the protein LoBind tubes (Eppendorf, Stevenage, UK) and kept at −20 °C for up to 2 weeks. Each day, an aliquot was thawed before administration to animals. Mice received a single subcutaneuos (s.c.) injection per day of 5 mg LiPR/kg of body weight before the onset of the dark cycle. As a control, LiPR vehicle was injected s.c.

### Liraglutide

Glucagon Like Peptide 1 (GPL-1) analogue Liraglutide (Saxenda, PubChem CID: 16134956, ^101^ was obtained from Pharmacy (Victoza®, Novo Nordisk A/S, Bagsværd, Denmark). Liraglutide was administered in the physiological solution (40 μg/ml) s.c. daily in 0.2 mg/kg concentration.

### Determination of hormonal and biochemical parameters

Blood was sampled from tail vein of overnight fasted animals and the blood plasma was separated and stored at −20°C. The plasma insulin concentrations were measured using an RIA assay (Millipore, St. Charles, MI, USA). Leptin concentrations were determined by ELISA (Millipore, St. Charles, MI, USA). The blood glucose levels were measured using a Glucocard glucometer (Arkray, Kyoto, Japan). The plasma triglyceride levels were measured using a quantitative enzymatic reaction (Sigma, St. Louis, MO, USA). The free fatty acids (FFA) levels were determined using a colorimetric assay (Roche, Mannheim, Germany). Cholesterol was determined by colorimetric assay (Erba Lachema, Brno, Czech Republic). All measurements were performed according to the manufacturer’s instructions.

### BrdU

Animals were given bromodeoxyuridine, BrdU (Sigma-Aldrich, Gillingham, UK) either by intraperitoneal (i.p.) injections or in drinking water. For i.p. injections, BrdU (10 mg/ml) was dissolved in the physiological solution (0.9% NaCl, pH = 7.4) and injected as 150 mg/kg of body weigh ^102^. Animals received 2 i.p. injections of BrdU 6 hours apart in the 7-day HFD protocol (see below). In drinking water, BrdU (1 mg/ml) was dissolved in sterile tap water with 1% sucrose ^61^. Animals were administered BrdU drinking water for 5 or 14 days (see the HFD protocols below). BrdU drinking water was protected against photolysis and changed every 3 days.

### HFD protocols

Mice were exposed to four different protocols. In the short protocol, they were given control or HFD with concurrent daily s.c. injections of LiPR or Liraglutide for 7 days and BrdU i.p. injections on day 1 and culled on day 7 (BrdU chase period of 6 days). In the intermediate protocol, they were exposed to 21 days of control or HFD with concurrent daily s.c. injections of LiPR or Liraglutide and administration of BrdU in the first 5 days and culled on day 21 (BrdU chase of 16 days). In the long protocol, mice were exposed to control or HFD for 4 months. On day 92, they were given BrdU drinking water for 14 days followed by daily s.c. LiPR injections for 14 days and culled on day 120 (BrdU chase of 14 days). In these 3 protocols, mice were culled by transcardial perfusion with brains used for immunohistochemistry. Finally, mice were exposed to 14 days of control or HFD with concurrent daily s.c. injections of LiPR and culled by cervical dislocation with brains used to prepare primary cell cultures or to isolate RNA for RNA sequencing. In the 14-day and 21-day protocols, LiPR was also administered to animals on Control (LFD) diet.

### Immunohistochemistry

Animals were transcardially perfused with ice-cold PBS followed by perfusion with ice-cold 4% paraformaldehyde. The brains were isolated and postfixed in 4% PFA for overnight. The PFA was replaced by 30% sucrose and brains let to sink at room temperature. 40μm coronal sections (spanning entire hippocampus) were cut in serial sets of 12 for stereological evaluation. Slide-mounted or free-floating immunohistochemistry (IHC) was performed. The following primary antibodies were used: rabbit polyclonal anti-GPR10 receptor (1:200; PA5-29809 Thermo Scientific, Paisley, UK); rabbit polyclonal anti-NPFF2 receptor (1:200; ab1420 AbCam, Cambridge, UK); chicken polyclonal anti-MAP2 (1:400; a92434 AbCam); mouse monoclonal anti-HuC/HuD (1:500; A21271 Thermo); mouse monoclonal anti-NeuN (1:500; MAB377 Thermo); mouse monoclonal anti-beta-III-Tubulin (1:400; ab14545 AbCam); rabbit polyclonal anti-GFAP (1:500; Z0334 Agilent/Dako, Stockport, UK); goat polyclonal anti-Vimentin (1:50; AB1620 Chemicon, Watford, UK); mouse monoclonal anti-Ki67 (1:400; CPCA-Ki67-100ul EnCor Biotechnology/2BScientific, Upper Heyford, UK); mouse monoclonal anti-PCNA (1:300; M087901-2 Agilent); guinea pig polyclonal anti-Doublecortin (DCX; 1:400; AB2253 Thermo); rat monoclonal anti-BrdU (1:400; MCA6144 BioRad, Watford, UK); guinea pig polyclonal anti-RAX (1:200; M229 Takara Bio Europe Ab, Goteborg, Sweden); rabbit polyclonal anti-activated caspase 3 (AC3; 1:500; 9661S Cell Signalling, Leiden, The Netherlands). Blocking with the carrier (1x PBS, 0.5% Triton X-100, 5% normal donkey serum or 2% bovine serum albumin) for a minimum of 20 minutes at room temperature (RT) was followed by incubation with the primary antibodies at RT overnight. After 3 washes, the secondary antibodies were incubated for 1.5 hours at RT. Sections were counterstained with DAPI (1:1000; Roche). Slides were mounted and cover-slipped using ProLong Diamond antifade mountant (Thermo). For BrdU staining, brain sections were incubated with primary antibodies for GFAP and HuC/HuD or NeuN and followed by their respective antibodies as described above. After that, the sections were post-fixed in 4% PFA in PBS for 15 minutes in RT and treated with 2N HCl (20 minutes in RT) followed by 0.1 M sodium tetraborate (pH = 8.5; 10 minutes in RT). After equilibration in PBS, primary antibody for BrdU was applied overnight in 4 °C, followed by 3 washes, and the secondary antibody.

### Stereologic and Proportional Cell Analyses

Quantification of marker-positive cells in the Medial Basal Hypothalamus (MBH) and the Subgranural Zone (SGZ) of the hippocampus was performed stereologically as described before ^102^. All coronal sections containing the hippocampus (in general, from bregma −0.6 to −4.0 mm) and MHB (in general, from bregma −1.2 to −2.3 mm, ^103^ from one well in the series of 12 were stained as described above as tanycytes are present in coordinates from bregma −1.3 to −2.5 mm ^104^. Z-stack images were obtained in the ZEN Blue software (Zeiss) using 20X or 40X apochromatic objectives on the Observer.Z1 Zeiss LSM780 confocal microscope by an observer blind to the genotype of the brains. Density of GPR10-positive (GPR10+) and NPFF2+ puncta (co-localized with Map2 and βIII-Tubulin neuronal cytoskeleton structures) was quantified in the Dorsal Medial Nucleus (DMN) of the hypothalamus and expressed per area of tissue quantified. BrdU+Map2+ neurons expressing GPR10 in the MBH (including the ME) were quantified as a proportion of all BrdU+Map2+ cells. A cell was recognized as BrdU+Map2+GPR10+ if it had Map2+ processes around or extending from a BrdU+ nucleus and there was co-localization of GPR10+ puncta with the BrdU-associated Map2+ processes. To obtain estimates of absolute number of marker-positive cells per volume of tissue, all cells positive for one or more cell markers (e.g. BrdU, NeuN and DAPI) were quantified in brain sections containing MBH. The Region of Interest (ROI) quantified included the MBH parenchyma with the Arcuate (Arc), DMN and Ventromedial (VMN) Nuclei and the Medial Eminence (ME). In the hippocampus, the ROI included SGZ only. The area of quantified ROI in 2-4 brain section was measured in the ZEN software and multiplied by the brain section thickness to obtain the ROI volume. The cell density was quantified as number of cells per mm cubic of tissue. Quantifications of tanycytes was done in sections containing MBH, including the number of GFAP+ α-tanycytes in the 3V wall and the proportion of GFAP+Vimentin+ tanycytes. The number of GFAP+ α-tanycytes which project from the 3V wall into the parenchyma were quantified. α-tanycytes were defined as being located in the lateral walls of 3V and having a cell process greater than two cell soma lengths and projected into MBH parenchyma. The number of GFAP+ α-tanycytes in 1 mm of MBH per section was calculated as (1/volume (mm^3^) x number of GFAP+α-tanycytes, to allow comparison between brain sections of different sizes. Density of GFAP signal from MBH parenchyma was quantified in orthogonal projections of confocal z-stack images using the Integrated Density function of the ImageJ software.

### Cell Cultures

Primary cell cultures were prepared from the dissected walls of 3V of adult C57BL/6j male mice exposed to control or HFD with or without concurrent LiPR administration for 14 days or from untreated mice. Cells were prepared as adherent primary cell cultures or as neurospheres. Isolated brain was washed 3 times in ice-cold HBSS with 10 mM HEPES. Dissection was carried out in the same medium as brainwashing. First, anterior, and posterior coronal cuts were made to isolate the hypothalamus in the rostro-caudal axis, corresponding to approximately bregma −1.0 to −4.0 mm. Then, a horizontal cut was made at the level of the thalamus to separate the cerebral cortex and to level the dorsal aspect. With the ventral aspect facing up, using microdissection surgical scissors, cuts parallel to the 3V wall were made to isolate the 3V walls. Finally, cust were made in the rostro-caudal axis to separate the segment containing the Hypothalamic Ventricular Zone (HVZ) corresponding approximately to bregma −1.0 to −2.5 mm. The dissected tissue was digested for 30 minutes in 37 °C with trypsin (3.5 mg/5 ml; Sigma) and hyaluronidase (3.5 mg/ 5 ml, Sigma) as described previously ^61^. After gradient centrifugation, the cells were resuspended in DMEM/F12 GlutaMAX medium (Thermo) supplemented with penicillin (100 units/ml), streptomycin (100 μ ml, Gibco) and B27 supplement (1:50, Thermo). For neurospheres essay, the medium was also supplemented with EGF and α (both 10 ng/ml; Gibco) every 3-4 days and primary neurospheres were counted and imaged on day 5 and 10 of the culture to measure their diameter. On day 10, neurospheres were collected for RNA isolation. For primary HVZ cultures, EGF and bFGF2 (both 10 ng/ml) was added once at plating of cells. 20.000 cells per 0.5 ml of medium per well (24-well plate) were plated in PDL-coated wells. Cells were allowed to adhere overnight in a cell incubator before they were subjected to time-lapse imaging. After imaging, cell identity was determined by immunocytochemistry (ICC) for proteins specific to different cell types as described in greater details in the following sections.

### Time-Lapse Imaging

Primary HVZ cells from HFD/LiPR treated mice were prepared as described above. Primary HVZ cells from untreated mice were exposed to LiPR in vitro for 8 days (1 μM added every day during the first 3 days of the culture). Alternatively, primary HVZ cells from naïve C57BL/6j mice were prepared and subjected to 1 μM hPrRP31 for 4 days (hPrRP31 added once per day) before the time-lapse imaging. Cells were continuously imaged as described previously ^61,105^. Briefly, cells were kept in 37 °C in 5% CO_2_ and pictures were taken every 10 minutes for 4 days. Four imaging regions (of 2 x 2 tiles) were selected per well in the ZEN Blue software (Zeiss). Imaging was done with an inverted 10X apochromat objective on a Zeiss Axio Observer 7 microscope equipped with an environmental chamber, motorized stage and Zeiss Definitive Focus module. After imaging, cells were stained for Vimentin, Sox2 and/or GFAP and their immunocytochemical phenotype was determined. Marker positive cells with elongated soma and one or two elongated processes were distinguished from rare astrocytes with stellate morphology. The cell dynamics of individual cell clones from a single seeding stem cell was analyzed either in ZEN software in less mitotically active clones (up to 8 cells per clone at the end of imaging) or in the The Tracking Tool for more active clones as described previously Timm’s Tracking Tool software as described previously ^61,106^.

### Immunocytochemistry

Cells were stained in 1X PBS with 2% bovine serum albumin (BSA, Sigma) and 0.5% Triton-X overnight in RT. After washout, the cells were stained with secondary antibodies (1:300) for 2 hours at room temperature and nuclei stained with DAPI (1:1000, Roche, Munich, Germany). The cells were mounted using the ProLong Diamond antifade mountant (Thermo) and imaged in the ROIs identical to the time-lapse ROIs.

### RT-qPCR

Total RNA was isolated from HVZ neurospheres using the RNeasy Mini plus kit (Qiagen, Manchester, UK) according to manufacturer’s manual. RNA was re-transcribed into cDNA using Super Script III polymerase with random primers and RNAse Out (Thermo). qPCR reaction was performed using Fast SYBR Green dye (Thermo) in technical duplicates for each sample on a Step One Plus Real time PCR system (Life Technologies) to determine relative expression of selected mRNA transcripts (denaturation for 5 min at 95 °C, followed by 40 cycles of 95 °C denaturation and 60 °C annealing and elongation). The results were analyzed using the AccuSEQ software (Life Technologies) with the amplification cycle (Ct) values determined by maximum 2^nd^ derivation method and following the 2^-ΔΔCt^ method ^107^ with normalization to the expression levels of Gapdh. The sequences of used primers (250 nM final concentration) were designed by using the Primer-BLAST online tool (NCBI-NIH): Gapdh forward = TTCACCACCATGGAGAAGG, Gapdh reverse = CACACCCATCACAAACATGG ^61^; Prlh forward = GCTGCTGCTAGGCTTAGTCC, Prlh reverse = ACTTGGCACTTCCATCCAGT ^108^; Prlhr forward = ACTTCCTCATTGGCAACCTG; Prlhr reverse = TGGTGAGTGTGAACACCGAT ^108^; Npffr2 forward = TGCCTATCACATTGCTGGAC; Npffr2 reverse = CGTGACAAAGGCTGTCTTGA ^108^; Vim forward = CACTAGCCGCAGCCTCTATTC, Vim reverse = GTCCACCGAGTCTTGAAGCA ^109^; Ttr forward = TCCTCATTTTTCTCCCCTGCT, Ttr reverse = CGGTTGGTCCACTCTGCTTT; Il31ra forward = CCAGAAGCTGCCATGTCGAA; Il31ra reverse = TCTCCAACTCGGTGTCCCAAC ^110^; Stxbp3 forward = CTGCGACAAAATACGGGCAG; Stxbp3 reverse = GGGGAACAATGGGAACACCA; Rap1b forward = GTGAATATAAGCTCGTCGTGC, Rap1b reverse = ACACTGCTGTGCATCTACTTC ^111^.

### RNA sequencing

Mice were exposed to Control diet or HFD for 14 days with or without the concurrent administration of LiPR (as described above). There were these mouse groups used for the RNAseq: Control (LFD) diet, HFD, LFD + LiPR, and HFD + LiPR. The weight of each mouse was recorded every 2 days. At the end of the 14 days all the mice were culled. Mice were culled by cervical dislocation and brains collected in ice-cold HBSS with 10 mM HEPES (Thermo). Coronal, 500 μm sections were cut on a vibratome (Leica) and MBH tissue was sampled with Reusable rapid biopsy punch kit with 1.0 mm in diameter (WPI, Hitchin, UK). Sampled tissue punches were flash frozen on dry ice and store at -80 °C until RNA isolation by RNeasy Plus mini kit (Qiagen). RNA quality checking was completed with the Qubit fluorometer system (Invitrogen) and assessed using RIN (RNA integrity number, maximum score is 10). All samples had RIN above 8.5, most above 9. RNA was re-transcribed to cDNA, tagmented and finally sequenced on the Illumina next-gen sequencer (75bp cartridges) following the Illumina user guide at at the Genomic Hub of School of Biosciences, Cardiff University. The RNA input concentrations for cDNA library preparation were normalized to 380 ng. cDNA libraries were carried out with 13 rounds of amplification.

### RNAseq data analysis

Raw reads were quality checked using FastQC ^112^ and quality filtered with fastp ^113^. Paired-end reads for each sample were mapped to the *Mus musculus* genome GRCm39 using STAR ^114^. Duplicates were marked using Picard ^115^ and alignments were quantified using subread ^116^. For each of the three pairwise comparisons (LFD vs HFD, Control Diet vs Control Diet + LiPR, HFD vs HFD + LiPR), differentially expressed genes were identified using the DESeq2 pipeline as implemented in SARTools ^117,118^. Genes were excluded from the analysis unless they had < 100 reads in at least one sample, as were all mitochondrial genes. Differentially expressed genes were defined as having an absolute log_2_ fold change >= 0.5 and an adjusted *P* value of < 0.05. *P* values were adjusted for multiple testing using the Benjamini-Hochberg method ^119^. Heatmaps were created using ComplexHeatmap ^120^ and GO term enrichment analysis was run using clusterProfiler and GO.db ^121,122^. Analysis was run using R v4.2.1 in RStudio ^123^(RStudio Team, 2016).

### hESCs-derived hypothalamic neuron differentiation

The maintenance and directed differentiation of embryonic stem cells to hypothalamic neurons were performed as previously described with slight modifications^65,66^. Specifically, differentiated HUES9 hESCs (Passage 12-18; RRID:CVCL_0057) were plated on 12-well iMatrix-511 (Cat.No. T303; Takara Bio Inc) coated plates at a density of approximately 3 x 10^6^ cells/well in Synaptojuice medium (adapted from ^124^. Hypothalamic cultures were maintained in Synaptojuice medium and replated onto iMatrix-511-coated on Ibidi 35 x 12 mm imaging dishes (Cat.No. 81156; Ibidi) at a density of approximately 2 x 10^6^ cells/well on days 33-40 post-differentiation. Calcium imaging experiments were then carried out on replated hypothalamic cultures at days 40+ post-differentiation.

### Calcium Imaging

Hanks’ Balanced Salt Solution Phenol Red free (HBSS, Cat. No. 14025092; ThermoFisher scientific) with synaptic blockers was used as extracellular bath solution. The final composition of the extracellular bath solution contained (in mM) NaCl 138, KCl 5.3, CaCl_2_ 1.26, MgCl_2_ 0.49, MgSO_4_ 0.4, KH_2_PO_4_, NaHCO_3_, Na_2_HPO_4_ 0.34, D-Glucose 5.5, DL-AP5 100 µM (Cat. No. ab120271; Abcam), Picrotoxin 50 µM (Cat. No. 1128; Tocris), CNQX 30 µM (Cat. No. ab120044; Abcam), and Strychnine 20 µM (Cat. No. ab120416; Abcam). Concentrated stock solutions of 1 mM hPrRP31 were prepared in H_2_O and stored at - 20 °C until the day of each experiment. Test concentration of 1 µM hPrRP31 were prepared in HBSS plus synaptic blockers immediately before each experiment. Differentiated stem cell-derived hypothalamic neurons were re-plated on ibidi 35 x 12 mm imaging dishes (Cat.No. 81156; Ibidi) and loaded with the Fluorescent Dye-Based Rhod-3 AM following the manufactures’ instructions (Rhod-3 Calcium Imaging Kit, Cat.No. R10145; ThermoFisher scientific). Loading, incubation, and wash buffers were prepared in HBSS solution in the presence of synaptic blockers. Imaging dishes were placed on an Olympus BX51WI Fixed Stage Upright Microscope (RRID:SCR_023069) and imaged using a 16-bit high-speed ORCA Flash4.0 LT plus digital sCMOS camera (RRID:SCR_021971). Neurons were identified by Rhod-3 AM fluorescence using an excitation wavelength of 560 nm and an emission wavelength of 600 nm. Images were taken using a 40X objective and acquired at a rate of one frame per second (100 ms and HCImage (RRID:SCR_015041) software for acquisition. Neurons were perfused continuously at room temperature with HBSS solution (3 mL/min) using a gravity-driven perfusion system for the entire length of the experiment and the fluorescence time course was recorded. The time course of each experiment was as follow: After a washout period of 10 min, cells were perfused with extracellular bath solution containing 1 μM hPrRP31 for 10 minutes. After second washout of a period of 10 minutes neuron were stimulated by 50 mM KCl for two minutes. A final washout of 10 minutes followed the application of KCl. Each fluorescence time course was exported to Microsoft Excel (RRID:SCR_016137), and the change in fluorescence intensity as a function of time was expressed as (F□−□*F*_0_)/*F*_0_) or Δ*F*/*F*_0_, where F is the total fluorescence and *F*_0_ is the baseline fluorescence. The extent of hPrRP31 response was evaluated by comparing the change in the area under the curve (AUC) of the fluorescence time-course before, during, and after the perfusion of hPrRP31.

### Statistical analysis

Numbers of biological and technical repeats are provided in the Figure Legends and in the Method Details. Datasets were analyzed with Microsoft Excel and GraphPad Prism. The statistical analysis has adhered to the following procedural algorithm. First, it was determined if the data are normally distributed by D’Agostino & Pearson’s omnibus normality test. If the data set is smaller than N = 5, the Kolmogorov-Smirnov normality test was used. Second, if data were not normally distributed or if this could not be reliably determined due to small number of replicates, non-parametric statistical tests were used such as Mann-Whitney for un-paired experiments, Wilcoxon matched-pairs signed rank test for paired experiments, or Kruskal-Wallis test with Dunn’s post-hoc test (for group comparison). If data are normally distributed, parametric tests were used. For simple comparison of two data groups, the un-paired two-tailed T-Test was used. For multiple factor or group comparison, OW or TW ANOVA with the Bonferroni post-hoc test was used. The contingency distribution of data is tested by either Chi-square or Fisher’s exact tests. For non-parametric tests, the data were presented as median ± interquartile range (IQR). The IQR was calculated as R = P * (n + 1)/100, where P is the desired percentile (25 or 75 for quartiles) and n is the number of values in the data set ^125^. For parametric tests, the data were presented as mean ± standard error of mean (SEM). Results were considered significant with P < 0.05 (one asterisk). In graphs, two asterisks represent values of P < 0.01, three asterisks for P < 0.001.

---

**Supplementary Figure 1.**
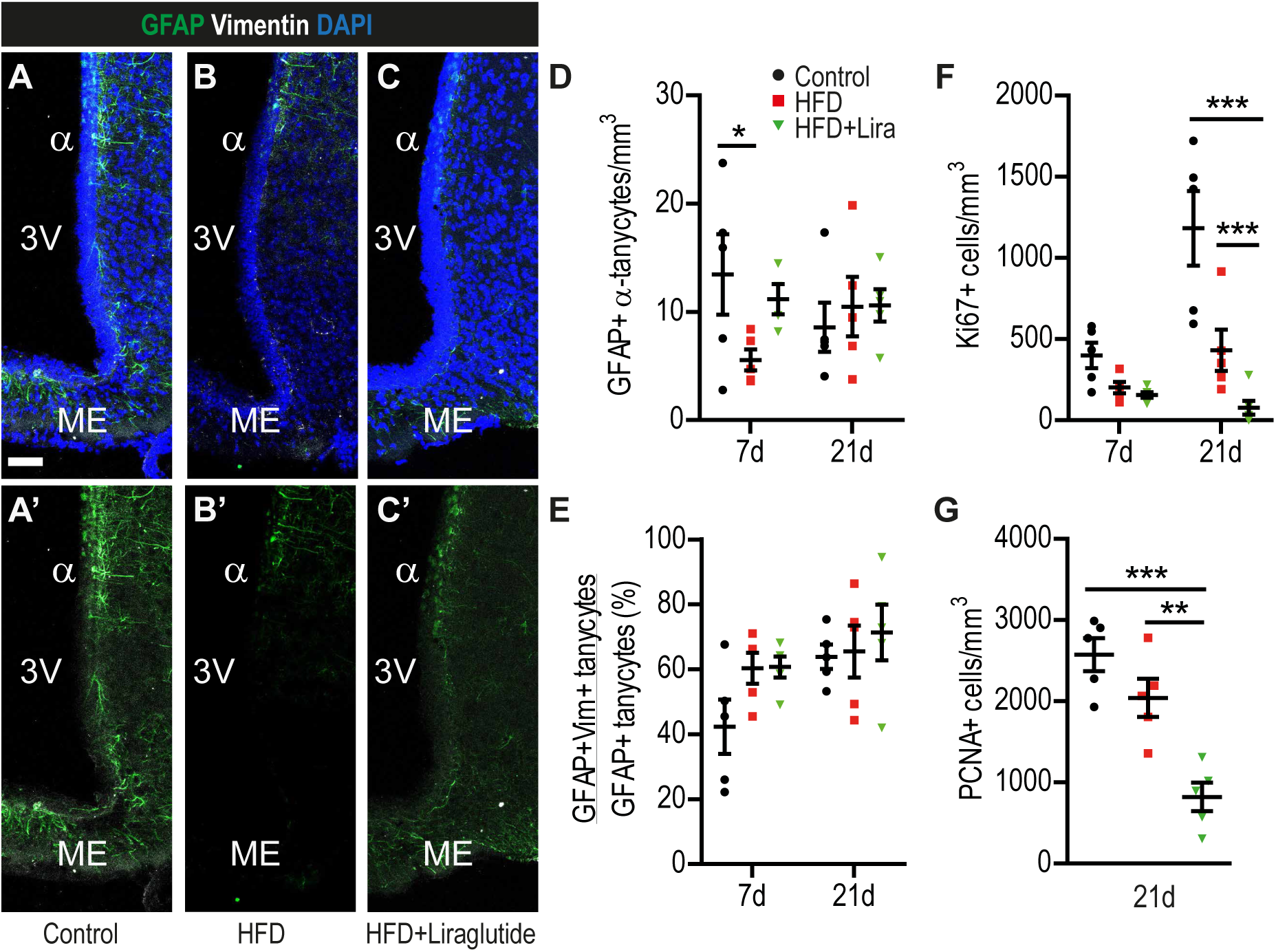
(A-C) Representative confocal images of HVZ stained as indicated in Control (A), HFD (B) and HFD + Liraglutide (Lira) (C) of 21d HFD group. (D) Quantification of GFAP+ α-tanycytes per volume of MBH. (E) Proportional analysis of GFAP+Vimentin+ tanycytes in MBH. (F-G) Quantification of MBH cells positive for Ki67 (F) and PCNA (G). Scale bars: 50 μm. n = 5, one-way ANOVA with Bonferroni’s test. *p < 0.05, **p < 0.01, ***p < 0.001. Data are presented as mean ± SEM.

**Supplementary Figure 2.**
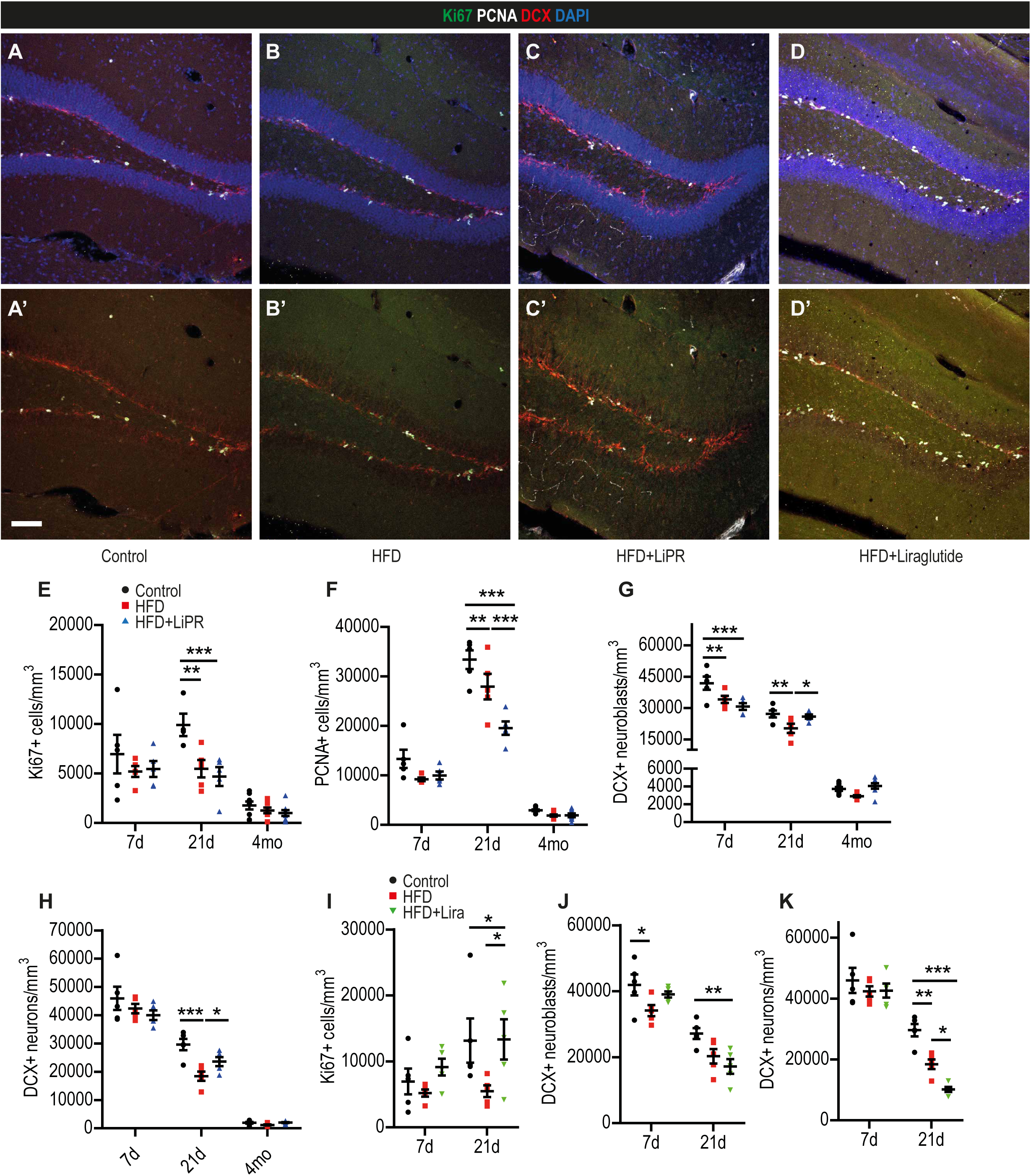
(A-D) Representative confocal images of the Dentate Gyrus (DG) of the hippocampus showing the Subgranular Zone (SGZ) stained as indicated in Control (A), HFD (B), HFD + LiPR (C) and HFD + Lira of 21d HFD group. (E-H) Quantification of Ki67+ (E), PCNA+ cells (F), DCX+ neuroblasts (G) and DCX+ neurons (H) in SGZ for Control, HFD and HFD + LiPR. (J-K) Cell quantification in SGZ as described for Control, HFD and HFD + Liraglutide. Scale bars: 50 μm. n = 5 for 7d and 21d groups, n = 8 for 4mo group. One-way ANOVA with Bonferroni’s test. *p < 0.05, **p < 0.01, ***p < 0.001. Data are presented as mean ± SEM.

**Supplementary Figure 3.**
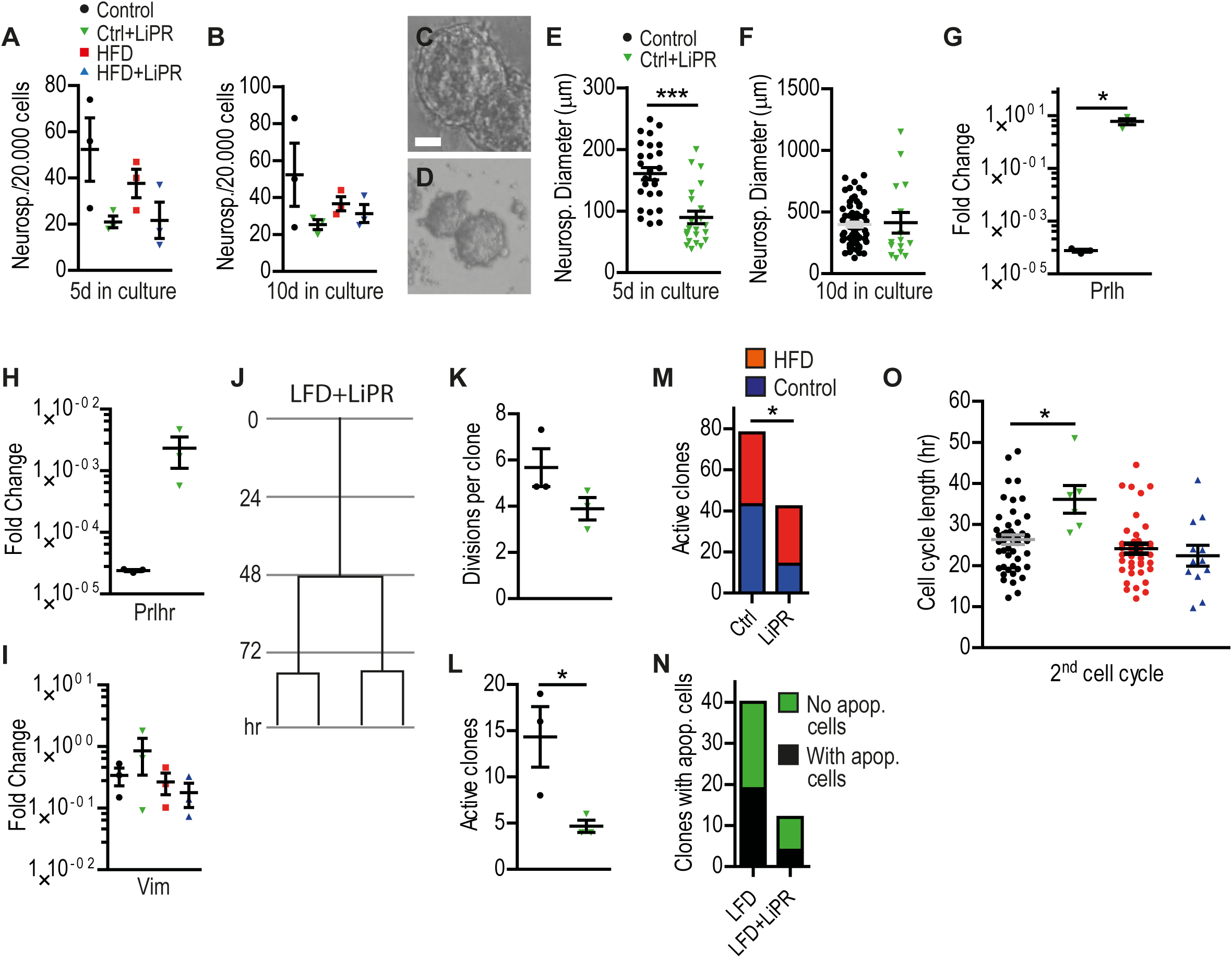
(A-B) Quantification of number of HVZ-derived neurospheres per 20.000 plated cells after 5d (A) and 10d (B) in culture. (C-D) Representative images of HVZ-derived neurospheres 5d in culture from Control (C) and Control + LiPR (D) treated mice. (E-F) Quantification of diameter of neurospheres 5d (E) and 10d (F) in culture. (G-I) Relative mRNA fold change of Prlh (G), Prlhr (H) and Vimentin (I) compared to Gapdh in cDNA from MBH-derived neurospheres (10d in culture). (J) An example cell division tree from 4-day time-lapse imaging of aNSCs from HVZ of Control + LiPR treated mouse. (K) Quantification of the number of cell divisions per division tree. (L) Time-lapse quantification of active (dividing) clones per 20.000 plated cells. (M) Number of all active clones of HVZ aNSCs from Control or LiPR-treated mice exposed to Control or HFD. Active clones pooled from all observed time-lapse imaging regions of interests per given treatment group. (N) Proportion of active clones containing at least one apoptotic cell. (O) Cell cycle length from the time-lapse imaging in the 2^nd^ observed cell division.Scale bar: 20 μm. n = 3. Fisher’s exact test (K, L), Un-paired two-tailed T-Test (E, F, G, H, K, L) and two-way ANOVA with Bonferroni’s test (A, B, O). *p < 0.05, ***p < 0.001. Data are presented as mean ± SEM.

**Supplementary Figure 4.**
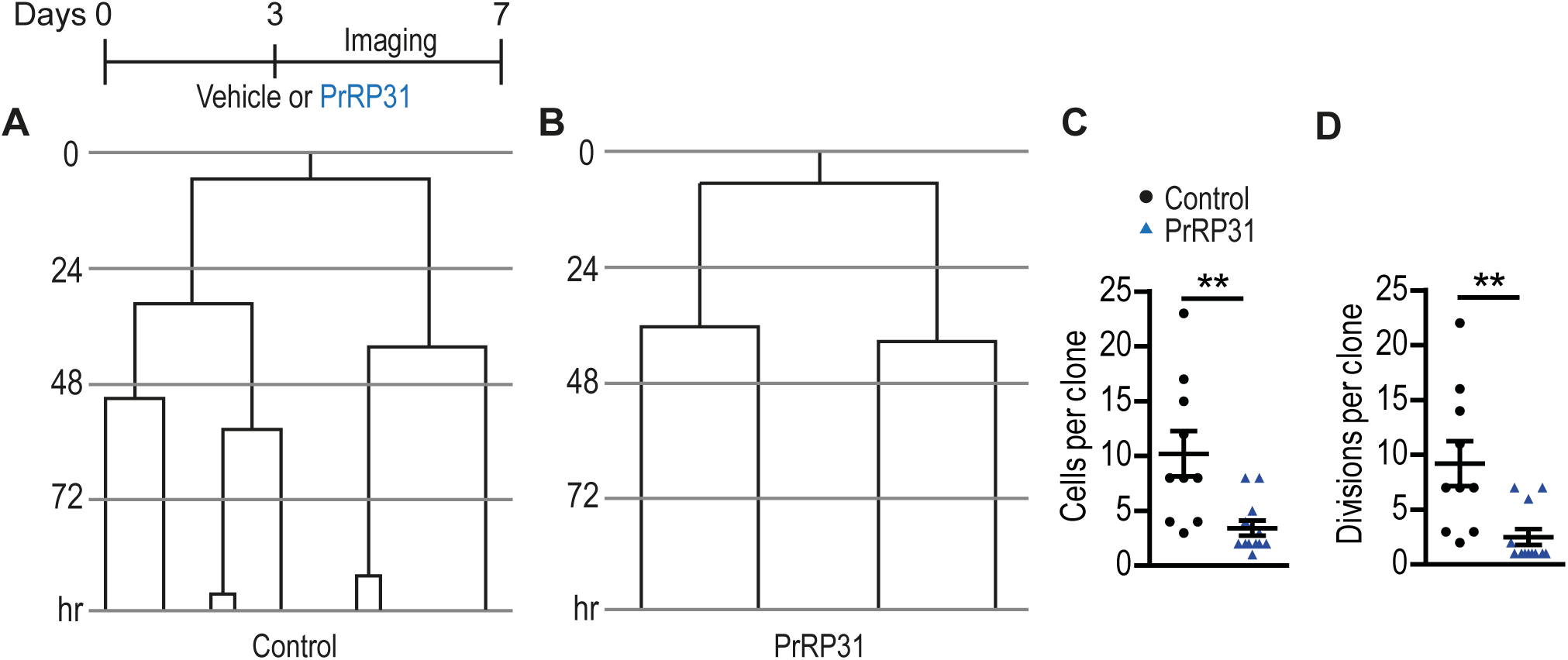
(A-B) Example cell division trees from 4-day time-lapse imaging of aNSCs from HVZ of Control (A) and hPrRP31-treated (B) cells. A schematic of the experimental protocol shown above. (C-D) Quantification of cells per clone (C) and divisions per clone (D). n = 3, un-paired two-tailed T-Test. **p < 0.01. Data are presented as mean ± SEM.

**Supplementary Figure 5.**
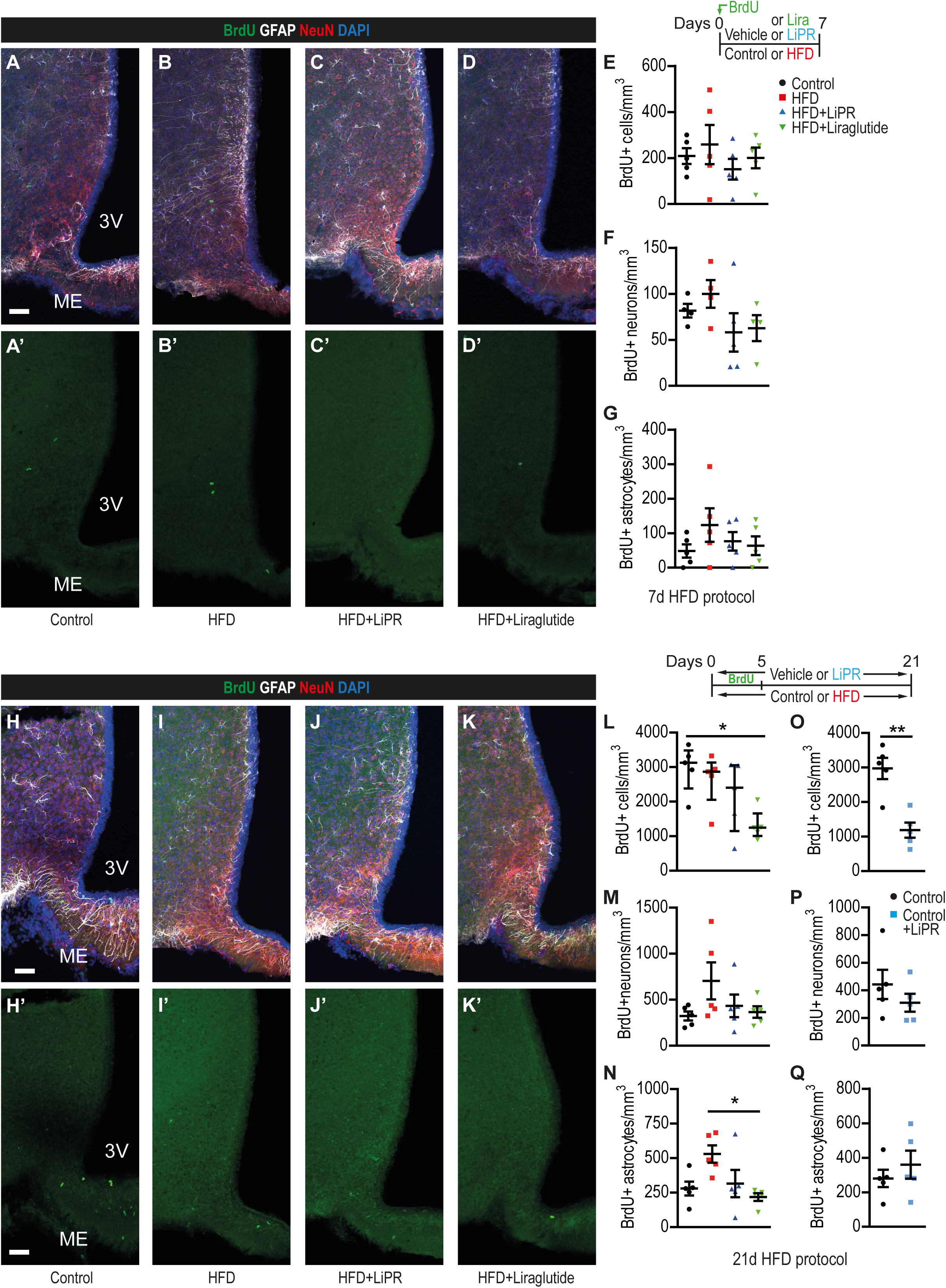
(A-D, H-K) Representative confocal images of HVZ stained as indicated in Control (A, H), HFD (B, I), HFD + LiPR (C, J) and HFD + Liraglutide (Lira) (D, K) of 7d (A-D) and 21d (H-K) HFD groups. (E-G) Quantification of BrdU+ cells (E), BrdU+ neurons (F) and BrdU+ astrocytes (G) in SGZ of 7d HFD group. (L-N) Quantification of BrdU+ cells (L), BrdU+ neurons (M) and BrdU+ astrocytes (N) in SGZ of 21d HFD group. (O-Q) Quantification of BrdU+ cells (O), BrdU+ neurons (P) and BrdU+ astrocytes (Q) in SGZ of Control and Control + LiPR treated mice of 21d group. Scale bars: 50 μm. n as in Fig.1. Un-paired, two-tailed T-Test (O-Q), Kruskal-Wallis test with Dunn’s test (L) or one-way ANOVA with Bonferroni’s test (all other). *p < 0.05, **p < 0.01. Data are presented as median ± SEM (L) or mean ± SEM (all other).

**Supplementary Figure 6.**
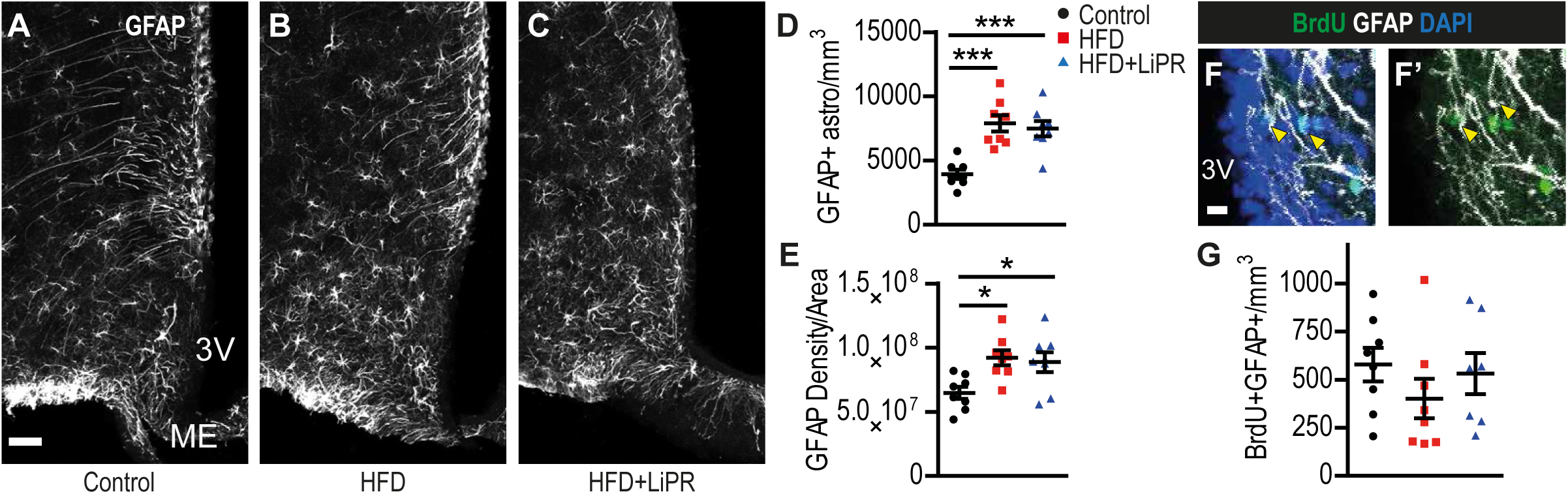
(A-C) Representative confocal images of HVZ stained for GFAP in Control (A), HFD (B) and HFD + LiPR (C) of 4mo HFD group. (D-E) Quantification of GFAP+ astrocytes (D) and GFAP density (E) in MBH parenchyma of 4mo HFD group. (F) Representative confocal image of BrdU+ astrocytes in MBH parenchyma. (G) Quantification of BrdU+ astrocytes in MBH parenchyma. Scale bars: 50 μm (A-C), 20 μm (F). n = 8, one-way ANOVA with Bonferroni’s test. *p < 0.05, ***p < 0.001. Data are presented as mean ± SEM.

**Supplementary Figure 7.**
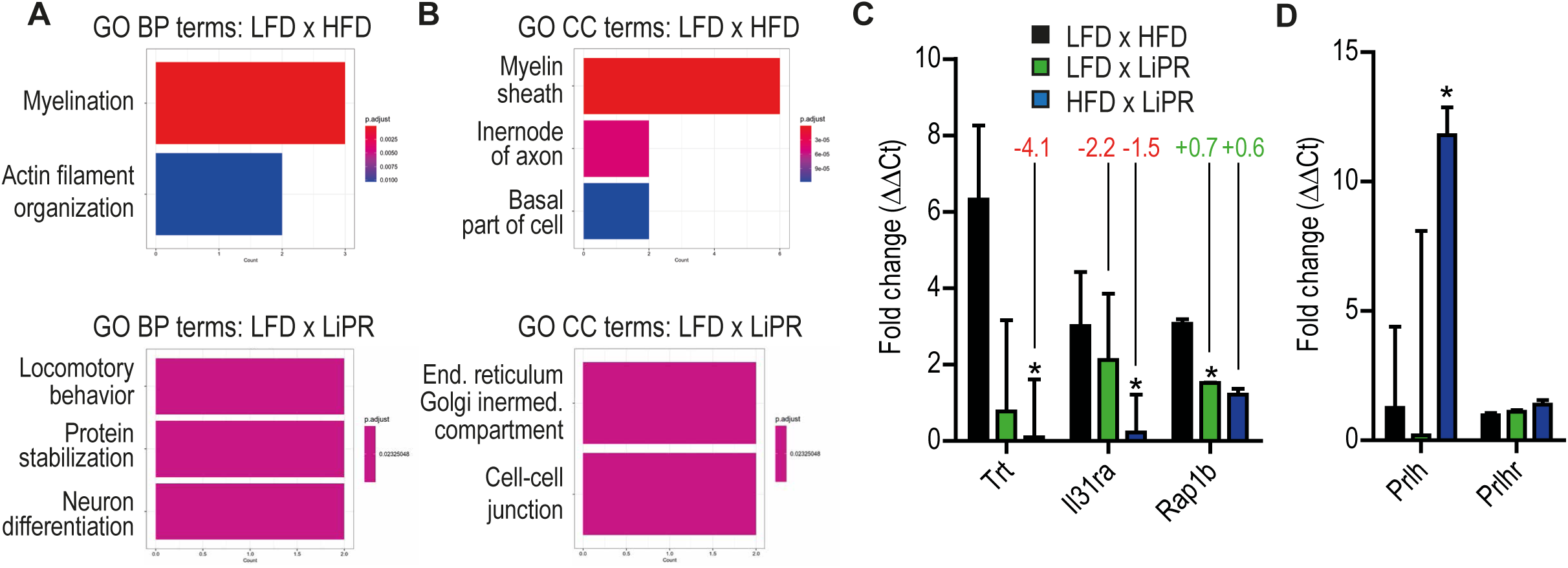
(A) Bulk MBH RNAseq GO terms for Biological Processes (BP) for LFD vs HFD (top graph) and LFD vs LFD+LiPR (bottom graph). (B) Bulk MBH RNAseq GO terms for Cell Compartments (CC). (C) Relative mRNA fold change of Trt, Il31ra and Rap1b compared to Gapdh in cDNA used for the bulk RNAseq of the MBH. Numbers above the bar graphs represent expression change values for a given gene from the RNAseq data. (D) Relative mRNA fold change of Prlh and Prlhr compared to Gapdh in cDNA used for the bulk RNAseq of the MBH. Un-paired two-tailed T-Test, *p < 0.05. Data are presented as mean ± SEM.

**Supplementary Figure 8.**
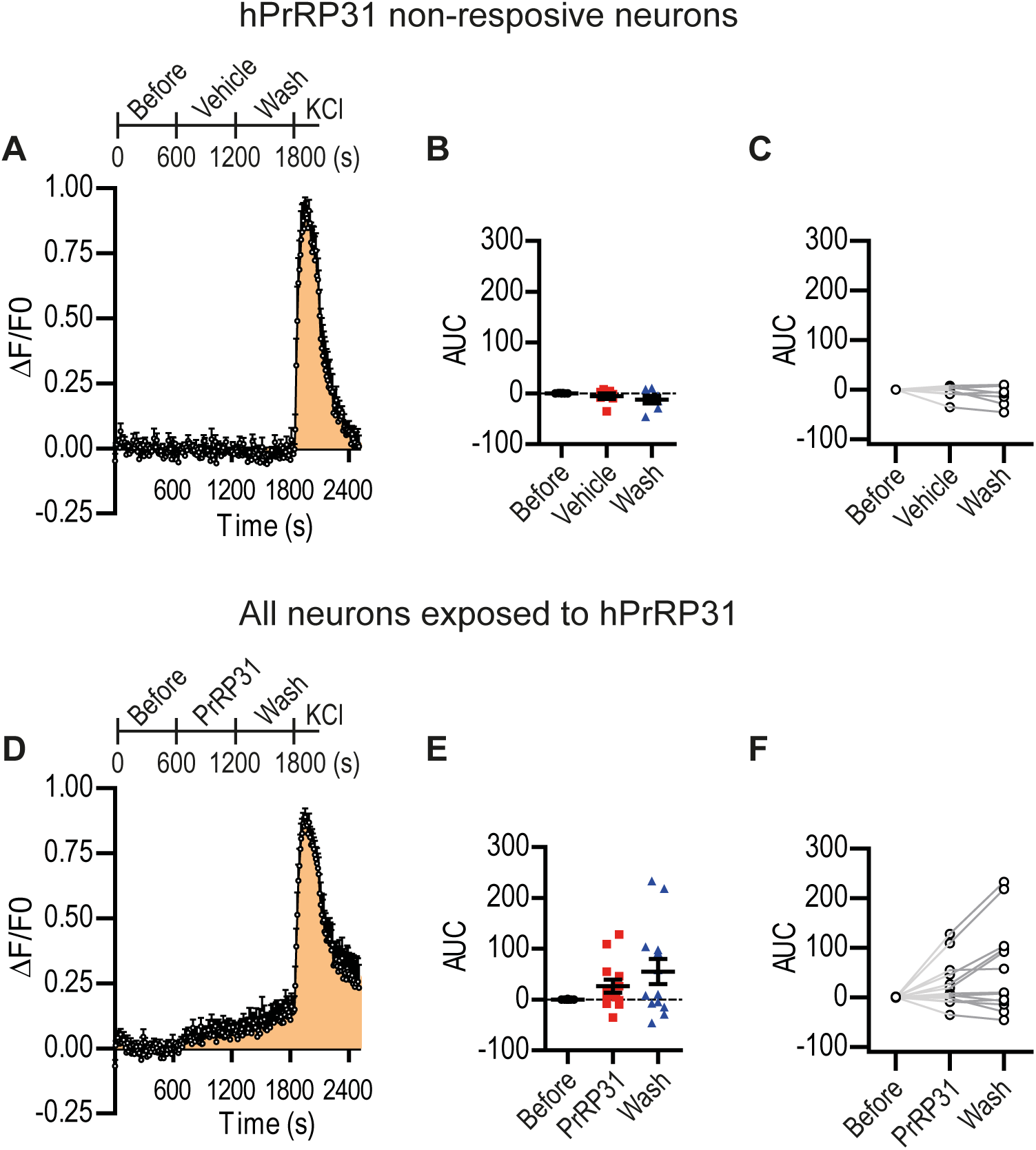
(A,D) A graph of the relative fluorescence change (ΔF/F0) of Rhod-3 as a function of time before and during hPrRP31 application in medium, wash-out and KCl positive control in hiPSC-derived neurons non-responsive to hPrRP31 (A) and all neurons exposed to hPrRP31. (B,E) A summary of fluorescence before (0-600 s), during hPrRP31 (600-1200 s) or wash- out (1200-1800 s) in Area Under Curve (AUC) for hPrRP31 non-responsive (B) and all neurons exposed to hPrRP31 (E). (C,F) Individual neurons from calcium imaging in the before-after plot from the non- responsive (C) and all exposed neurons (F). n = 7 for non-responsive, n =13 all neurons. Data are presented as mean ± SEM.

